# Cognitive maps for hierarchical spaces in the human brain

**DOI:** 10.1101/2025.02.05.636580

**Authors:** Michael Peer, Russell A. Epstein

## Abstract

Many of the environments that we navigate through every day are hierarchically organized—they consist of spaces nested within other spaces. How do our mind/brains represent such environments? To address this question, we familiarized participants with a virtual environment consisting of a building within a courtyard, with objects distributed throughout the courtyard and building interior. We then scanned them with fMRI while they performed a memory task that required them to think about spatial relationships within and across the subspaces. Behavioral responses were less accurate and response times were longer on trials requiring integration across the subspaces compared to trials not requiring integration. fMRI response differences between integration and non-integration trials were observed in scene-responsive and medial temporal lobe brain regions, which were correlated the behavioral integration effects in retrosplenial complex, occipital place area, and hippocampus. Multivoxel pattern analyses provided additional evidence for representations in these brain regions that reflected the hierarchical organization of the environment. These results indicate that people form cognitive maps of nested spaces by dividing them into subspaces and using an active cognitive process to integrate the subspaces. Similar mechanisms might be used to support hierarchical coding in memory more broadly.

## INTRODUCTION

Our navigable world consists of spaces at different scales: rooms, buildings, neighborhoods, cities, countries. These spaces are often hierarchically nested within each other—rooms are contained within buildings, which in turn are contained within neighborhoods. How do our minds/brains represent this hierarchical organization? And how do they mediate the relationship between different levels of the hierarchy, to allow us to navigate from place to place?

One possibility is that our minds create spatial representations of the environment (“cognitive maps”) that are themselves hierarchically organized. Previous work in cognitive psychology provides evidence for this idea: we mentally divide large environments into subspaces and represent the spatial relationships between subspaces separately from the spatial relationships within subspaces. Evidence for hierarchical coding is observed in geographical spaces (Stevens & Coupe, 1978), real-world navigable spaces (Hirtle & Jonides, 1985; McNamara, 1986), and 2-d figural spaces (Huttenlocher et al., 1991). It is observed across multiple behavioral paradigms, including directional judgements (Stevens & Coupe, 1978), position estimates (Huttenlocher et al., 1991), distance estimates (Hirtle & Jonides, 1985), free recall (Hirtle & Jonides, 1985; Taylor & Tversky, 1992) and navigational planning (Wiener et al., 2004; Wiener & Mallot, 2003). It is observed when subspaces are physically bounded by walls and barriers (McNamara, 1986) and when subspace divisions are mentally imposed top-down in the absence of any physical boundaries (Huttenlocher et al., 1991; McNamara, 1986).

Some of the most compelling evidence for hierarchical coding comes from the fact that people find integration across levels of the hierarchy to be challenging. For example, in one study, participants found it very difficult to point to campus landmarks while inside a campus building, even though they could do so accurately if they exited the building, thus putting themselves in the same hierarchical level as the landmarks (Wang & Brockmole, 2003a). In another study, a switching cost was observed when participants were required to recall spatial relationships at different hierarchical levels (room, building, campus) in successive trials ((Brockmole & Wang, 2002); see also (Wang & Brockmole, 2003b)). Results such as these suggest the existence of separate spatial representations for different levels of the hierarchy, which can only be integrated with an active mental process that is sometimes prone to failure. Integration across levels may be particularly difficult when the levels have principal axes that are angularly offset from each other; when this happens, people can develop different reference frames for each level, as evidence by alignment effects in judgment of relative direction tasks (Adamou et al., 2014; Greenauer & Waller, 2010; Kelly et al., 2018; Meilinger et al., 2014; Strickrodt et al., 2019; H. Zhang et al., 2014).

Despite this extensive behavioral evidence, little is known about how hierarchical spaces are encoded in the brain, or how the brain integrates across different levels of a spatial hierarchy. The majority of studies in spatial neuroscience have little to say about this issue, because they examine brain responses in environments that have a single uniform scale and/or a single space (e.g., a single room, or a continuous city region). A few studies have looked at where different scales of space are represented by examining memory recall in differently-sized spaces (Peer et al., 2019), the intrinsic dynamics of fMRI responses during navigation (Brunec et al., 2018; Brunec & Momennejad, 2022), or activity during navigation related to use of local vs. global spatial strategies (Evensmoen et al., 2013). These studies have demonstrated gradients of responses in the hippocampus, scene-selective regions, and prefrontal cortex related to scale, with more posterior regions responding to smaller spatial scales and more anterior regions responding to larger scales (Brunec et al., 2018; Brunec & Momennejad, 2022; Evensmoen et al., 2013; Peer et al., 2019). However, these studies did not examine how these spaces at different scales are represented, or how the scales are integrated with each other.

Other fMRI studies have used multivoxel pattern analysis and adaptation analyses to explore how the brain represents environments that are divided into subspaces. These studies have revealed evidence for both “local” (within-subspace) and “global” (across-subspace) spatial codes. For example, “spatial schemas” representing the common local coordinate frame shared by geometrically similar subspaces have been observed in the hippocampus (Kim & Maguire, 2018; Peer & Epstein, 2021), retrosplenial/medial parietal region (Marchette et al., 2014) and the scene-selective occipital place area (Peer & Epstein, 2021), whereas global coordinate frames that span multiple subspaces have been observed in the retrosplenial/medial parietal region (Peer & Epstein, 2021). Heading codes anchored to the local reference frame have been observed in the retrosplenial/medial parietal region (Marchette et al., 2014; Shine & Wolbers, 2021), whereas heading codes anchored to the global reference frame have been identified in the thalamus (Shine et al., 2016). To our knowledge, however, no study has looked specifically at how our brains bridge across hierarchical levels at different spatial scales.

In the current experiment, we addressed this issue by training participants on a hierarchically organized environment consisting of two nested subspaces – a building inside a courtyard. The two subspaces were visually distinct from each other and their main axes were rotated with respect to each other, thus encouraging the creation of separate local maps with different reference frames. Following learning, participants were scanned with fMRI while performing a judgment of relative direction task that required them to retrieve spatial relations within each subspace and across the two subspaces, thus allowing us to study the brain mechanisms involved in subspace integration. To anticipate, our findings show that people form separate cognitive maps of the two hierarchy levels (the building and its surrounding courtyard), use the hierarchical structure of the environment to make inferences about the relation between the levels, and integrate across the subspaces by using an active cognitive process mediated by scene-responsive and medial temporal lobe brain regions.

## MATERIALS AND METHODS

### Participants

32 healthy individuals (10 male, 21 female, 1 undisclosed sex; mean ± SD age 24.3 ± 4.4 y) from the University of Pennsylvania community participated in the experiment. This number was selected based on a power analysis on similar effects in our previous study which used similar methods (Peer & Epstein, 2021). All participants had normal or corrected-to-normal vision and provided written informed consent in compliance with procedures approved by the University of Pennsylvania Institutional Review Board (approval ID 843764). Three additional participants started the experiment but were excluded before analysis: two for failing to complete the spatial learning task in the allotted time on day 1, and one for failing to arrive for the day 2 scan.

### Experimental Design

#### Virtual environment

We used Unity 3D software to construct a hierarchically organized virtual environment consisting of a smaller space (a building) nested within a larger space (a courtyard). The courtyard was 300 x 400 virtual meters, and its boundaries were defined by impenetrable barriers – cliffs on its two long sides (east and west), a snow-capped mountain at its northern end, and a forest at its southern end. The building was 75 x 100 virtual meters, and its interior had brick walls, a ceiling, a concrete stripe on the floor extending from the door to the far (east) wall, and three columns in the center of the east wall. The courtyard and the building interior were connected by a single doorway on the “west” side of the building (Fig. 1A). The main (long) axes of the two spaces were perpendicular to each other, and both spaces contained elements that emphasized these axes (mountain and forest for the courtyard, door-stripe-columns for the buildings).

**Figure 1.**
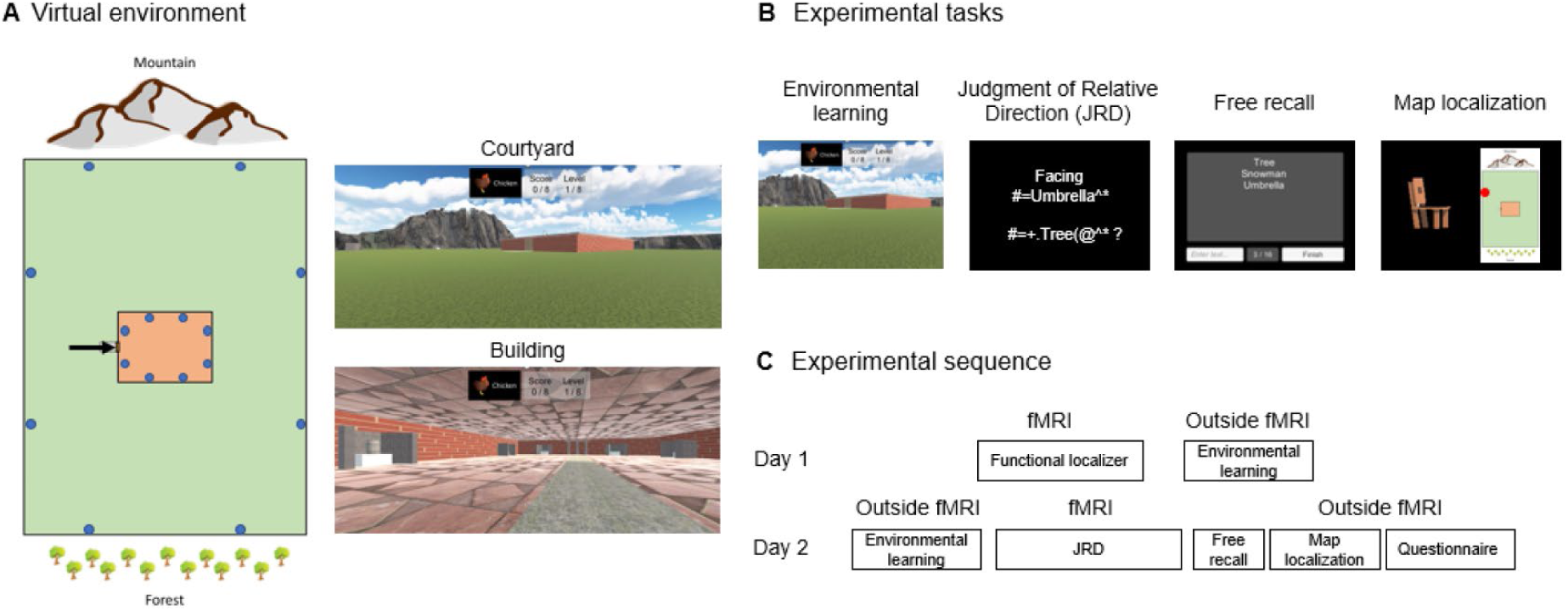
Experimental design and procedure. A) Participants were familiarized with a virtual environment consisting of a building (orange) inside a courtyard (green). The building interior could be accessed through a single doorway on the “West” side (marked by an arrow in the image). The building interior could not be perceived from the courtyard or vice versa. Sixteen objects (marked in the image by blue dots) were located in the environment, eight in each subspace (building and courtyard). Participants only saw the environment from a ground-level perspective (right) and never saw an overhead map (left). B) Experimental tasks. Images show the visual displays presented to the participants during the experimental tasks. C) Experimental procedure. Participants learned the locations of the objects in the environment (Environmental Learning) outside the fMRI scanner on Day 1, with a refresher on Day 2. Then the Judgment of Relative Direction (JRD) task was performed within the scanner, followed by free recall and map localization tasks outside the scanner. See the Methods section for full details on the tasks and procedure.

Sixteen objects were located within the environment, eight along the boundaries of the courtyard (two along each boundary) and eight inside the building along its interior walls (two along each wall). The objects were located on white pedestals that were surrounded on three sides by walls forming an alcove, so that they could only be viewed and approached from one direction (i.e., facing the boundary against which they were situated). The sixteen objects were: pyramid, chicken, statue of a person, umbrella, basket, motorcycle, barrel, traffic cone, guitar, treasure chest, hammer, mushroom, banana, chair, tree, and snowman. The assignment of these objects to the sixteen locations was randomized across participants.

#### Experimental sequence

The experiment consisted of two sessions, performed on consecutive days (except for one participant who had a four-day gap between scans due to MRI failure in the second day). Behavioral and MRI data were collected on both days (Fig. 1b and 1c). This subsection describes the sequence of tasks, while the following subsections describe each task in detail.

On day 1, participants were familiarized with the objects used in the experiment (Object Familiarization Task). They were then scanned with fMRI. The MRI scanning included functional localizer scans, anatomical T1 acquisition scans, an object viewing task in which participants viewed the objects and made simple perceptual judgments, and a resting-state scan in which participants were instructed to keep their eyes open and make no response. (The object viewing and resting-state data were not analyzed in this report; hence these tasks are not described further.) After exiting the scanner, participants performed the Environmental Learning Task, which provided intensive training on the spatial layout of the environment and the locations of the objects within it. This was the first time they saw the objects in the context of the virtual environment. Finally, they were given brief initial training on the Judgment of Relative Direction (JRD) task, in preparation for using the task the next day (data from these day 1 training trials were not analyzed). All told, session 1 lasted on average 80 minutes: 50 min for the fMRI session, 27 min for the post-scan environmental training and 3 min for the JRD training.

On day 2, participants performed ten additional minutes of the Environmental Learning Task to refresh their memory of the spatial layout of the virtual environment. They then entered the MRI scanner and performed two runs of the JRD task, which served to elicit MRI activity corresponding to explicit retrieval of spatial information. The day 2 MRI session also included additional runs of the object viewing task. They then exited the scanner and performed the Free Recall and a Map Localization Tasks, to further query their spatial memories for the virtual environment, followed by a post-experiment questionnaire. All told, session 2 lasted approximately 80 minutes (45 min for the fMRI session and 35 min for the behavioral tests).

Experimental tasks were programmed in Unity 3D and Psychopy 3 (Peirce, 2007), and stimuli sequences were selected using custom MATLAB code and using the webseq tool at https://cfn.upenn.edu/aguirre/webseq/ for carryover sequences (Aguirre, 2007).

#### Object familiarization task

To familiarize participants with the experimental objects they were shown each of the sixteen objects in random order on a black background. Participants were instructed to pay attention to the object images and names.

#### Functional localizer task

To enable identification of scene-selective brain regions, participants performed a 1-back repetition detection task while viewing 16 s blocks of faces, scenes, objects and scrambled objects, with each stimulus presented for 600ms followed by 400ms of fixation.

#### Environmental learning task

This task was intended to teach participants the structure of the environment and the locations of the objects within it. Participants freely navigated the virtual environment from a first-person perspective at a constant speed, using button presses to indicate their direction of movement. On each trial, they were given the name and image of an object and navigated to its location. The task was divided into eight learning stages of increasing difficulty, where the first six stages focused on the objects in one subspace (either the building or the courtyard – eight objects per learning stage), while stages 7-8 included all sixteen objects across the whole environment.

In stages 1 and 2, participants were instructed to look for eight objects in one of the subspaces (either the courtyard or building), while all objects were completely visible, in order to learn their locations. Stages 3-4 were similar, but in each trial four out of the eight objects were covered by a wooden box, so that these objects were not visible. The objects that were covered varied randomly from trial to trial, but always included the goal object; therefore, participants had to identify the goal object’s location from memory. In stages 5-8, all objects were covered. The order of the goal objects was randomized with each stage, and the order of subspaces in each pair of stages was randomized across participants. Participants started each stage in the center of the southern environment boundary, with their backs to the forest and facing the building (global “north”).

Each environmental learning stage began with short instructions, which indicated the location to search (building, courtyard, or the whole environment). The name and image of the first object was shown at the top of the screen. Participants were required to navigate to a position just facing the pedestal supporting this object and press the ‘‘down’’ key. If they selected the correct pedestal, a green light appeared, and the next goal object was indicated. If they selected an incorrect pedestal, a red light appeared, and the occluding box around the object was briefly removed to enable learning from the error. The trial then continued until the correct pedestal was found. After all relevant objects were found (eight objects in stages 1-6, sixteen objects in stages 7-8), participants were re-tested on any objects that they had made errors on, until they had found each object at least once without making an error; only then did they pass to the next stage. In stage 8, all objects had to be found in sequence while making a maximum of one error; if more than one error was made, the whole stage started anew. A counter at the top of the screen indicated how many objects had been found successfully during the current stage.

We established a pre-set limit of one hour overall to complete the task. Participants who exceeded this time limit were excluded from the rest of the experiment. The gradual learning, repetition of incorrectly remembered objects at the end of each stage, and requirement for near-perfect object-finding at the last stage were intended to ensure that participants accurately encoded all the object locations. Participants performed the full learning task to completion on day 1. On day 2, they performed 10 minutes of the task to refresh their memory of the environment, starting from stage 1. See Video S1 for a short demonstration of the environmental learning task.

#### Judgment of Relative Direction (JRD) task

Subsequent to environmental learning, participants performed a judgement of relative direction task, which was intended to elicit representations of spatial locations with the environment. On each trial, participants were presented with the names of two objects. They were instructed to imagine that they were standing facing the first object (starting object), and to indicate by button press whether the second object (target object) would be to the left or to the right given this imagined view. Each trial lasted 5 s and the names of the objects remained on the screen the entire time. Trials were followed by a variable inter-stimulus interval of 1 s (3/8 of the trials), 3 s (3/8 of the trials), or 5 s (1/4 of the trials). Object names were padded with non-letter characters to eliminate any fMRI response differences related to the number of letters in each name.

There were two consecutive JRD runs within the fMRI session, each lasting 516 s, with 64 unique trials per scan run. Trials could be of four trial types – 1) both objects were in the building; 2) both objects were in the courtyard; 3) the starting object was in the building and the target object was in the courtyard; 4) the starting object was in the courtyard and the target object was in the building. Trials with the starting object and the target object in different subspaces were labeled integration trials, because they required participants to spatially integrate across the subspaces. Trials with the starting and target object in the same subspace were labeled non-integration trials.

Sets of trials were constructed that fulfilled the following criteria: 1) there were an equal number of trials of each of the four trial types in each scan run (16 trials per trial type); 2) For each trial type, each object was used exactly two times per run as a starting object and two times as a target object; 3) The target object was never directly behind the starting object; 4) No trial was repeated throughout the experiment. Trials sets were randomly generated for each participant that met these four criteria. Trial sequences were then shuffled within each run, while controlling the order such that consecutive appearances of each of the four trial types (e.g., a trial of trial type 3 followed by trial type 1) occurred either 7 or 8 times, and consecutive trials never had the same facing object. These constraints on the transitions between trial types were intended to facilitate investigation of the switching costs related to subspace change between trials. During day 1 of the experiment, a training version of fifteen self-paced JRD trials followed by fifteen timed trials was performed outside the scanner, using object combinations that were not used in the main JRD task, which was performed in the scanner on day 2.

#### Free recall task

This task, performed at the end of the experiment, tested whether participants’ recall order was shaped by their subspace membership (Peer & Epstein, 2021). Participants were instructed to type the name of each object they could recall in the environment, in any order. They were instructed to press return after entering each word and press a ‘‘finish’’ button when they had written down as many names as they could recall. Participants saw all the words they had already entered on the screen (on different lines), as well as a counter indicating the number of words they have entered, but they could not go back and erase previously entered words. The task was self-paced.

#### Map localization task

This task, also performed at the end of the experiment, measured participants’ explicit knowledge of the locations of objects in the environment. Participants saw a map of the environment showing the boundaries of the courtyard, the walls of the building, the mountain, the forest, and the building door. The locations of the objects (i.e. the pedestals) and the alcoves were *not* indicated. On each trial, they were presented with the image and name of one object. They were instructed to click the cursor within the map to indicate the location of the indicated item, at which point a red dot appeared in the clicked location. Participants could click again to reselect the location as many times as they wished before finalizing their answer by clicking a “continue” button. The dot then disappeared, and the next trial began. Each of the sixteen objects was queried once, in random order. The task was self-paced.

#### Post-experiment questionnaire

Participants were asked to write down their strategy for solving each task, rate the difficulty of each task, estimate the environment’s size, and rate on a scale of 1-10 for each subspace (the courtyard and the building): how well they felt they knew the environment, how much they used first-person (eye-level) imagination during the JRD questions, and how much they used a third-person (bird’s-eye, map-like) imagination of the environment. Finally, they were asked to describe whether they memorized and thought of object locations relative to any landmarks (e.g. mountain, forest, building door/walls).

### MRI Acquisition and Preprocessing

Scanning was performed at the Center for Functional Neuroimaging at the University of Pennsylvania on a Siemens 3.0 T Prisma scanner using a 64-channel head coil. T1-weighted images for anatomical localization were acquired using an MPRAGE protocol [repetition time (TR) = 2,200 ms, echo time (TE) =

4.67 ms, flip angle = 8^0^, matrix size = 192 X 256 X 160, voxel size = 1 X 1 X 1 mm]. Functional T2*-weighted images sensitive to blood oxygen level dependent contrasts were acquired using a gradient echo planar imaging (EPI – EPFID) sequence (TR = 2,000 ms, TE = 25 ms, flip angle = 70^0^, matrix size = 96 X 96 X 81, voxel size = 2 X 2 X 2 mm).

Preprocessing was performed using fMRIPrep 20.2.6 (Esteban et al., 2019) (RRID:SCR_016216), which is based on Nipype 1.7.0 (Gorgolewski et al., 2011) (RRID:SCR_002502). The T1-weighted (T1w) image was corrected for intensity non-uniformity (INU) using N4BiasFieldCorrection (ANTs 2.3.3) (Tustison et al., 2010) (RRID:SCR_004757), and used as T1w-reference throughout the workflow. The T1w-reference was then skull-stripped using the antsBrainExtraction.sh workflow (ANTs 2.3.3), using OASIS30ANTs as target template. Brain parcellations into anatomical regions were defined using recon-all (FreeSurfer 6.0.1, RRID:SCR_001847) (Dale et al., 1999). Volume-based spatial normalization to MNI152NLin2009cAsym space was performed through nonlinear registration with antsRegistration (ANTs 2.3.3), using brain-extracted versions of both T1w reference and the T1w template (Fonov et al., 2009). Brain tissue segmentation of cerebrospinal fluid (CSF), white-matter (WM) and gray-matter (GM) was performed on the brain-extracted T1w using the fast tool (FSL 5.0.9, RRID:SCR_002823) (Y. Zhang et al., 2001).

For functional T2*-weighted scan runs, the following preprocessing was performed. First, a reference volume and its skull-stripped version were generated using a custom methodology of fMRIPrep. A B0-nonuniformity map (or fieldmap) was estimated based on a phase-difference map calculated with a dual-echo GRE (gradient-recall echo) sequence, processed with a custom workflow of SDCFlows inspired by the epidewarp.fsl script and further improvements in HCP Pipelines (Glasser et al., 2013). The fieldmap was then co-registered to the target EPI (echo-planar imaging) reference run and converted to a displacements field map (amenable to registration tools such as ANTs) with FSL’s fugue and other SDCflows tools. Based on the estimated susceptibility distortion, a corrected EPI (echo-planar imaging) reference was calculated for a more accurate co-registration with the anatomical reference. The T2* reference was then co-registered to the T1w reference using bbregister (FreeSurfer) which implements boundary-based registration (Greve & Fischl, 2009) using six degrees of freedom. Head-motion was estimated using mcflirt (FSL 5.0.9) (Jenkinson et al., 2002). Gridded (volumetric) resamplings were performed using antsApplyTransforms (ANTs), configured with Lanczos interpolation to minimize the smoothing effects of other kernels (Lanczos, 1964). Functional runs were slice-time corrected to using 3dTshift from AFNI 20160207 (Cox & Hyde, 1997) (RRID:SCR_005927) and resampled to MNI152NLin2009cAsym space. To measure potential confound variables, head motion framewise displacement (FD) was calculated for each preprocessed functional run, using its implementation in Nipype (following the definitions by (Power et al., 2014)); average signals were also extracted from the CSF and the white matter. Finally, functional data runs were smoothed with a Gaussian kernel (3mm full width at half maximum) using SPM12 (Wellcome Trust Centre for Neuroimaging).

### Behavioral analyses

#### Environmental learning task

We examined behavior in this task for evidence of segmentation between the subspaces. To do this, we compared within-subspace errors (choosing an incorrect location, but in the correct subspace) to between-subspace errors (choosing a location in the incorrect subspace) using a two-tailed paired-samples t-test. Trials were counted as correct if the first pedestal that participants selected was the location of the searched-for object. If it was not, the trial was counted as an error, even though the trial would continue until the correct location was found. For this segmentation analysis, we only considered responses in stages 7 and 8, as these were the stages where participants were required to search for object locations across both subspaces. We also calculated overall accuracy (the number of correct trials divided by the number of trials) in stages 3-8 to verify the learning curve, and to compare performance between courtyard and building objects. Performance in stages 1 and 2 was not assessed, because all objects were visible in those stages, including the target objects.

#### Judgment of Relative Direction (JRD) task

We used a 2 x 2 repeated-measures ANOVA (with participant as the random effect) to test for integration and subspace effects in the accuracy and reaction time data. Integration effects were defined as differences between trials requiring integration (starting and target objects in different subspaces) and trials not requiring integration (starting and target objects in the same subspace). Subspace effects were defined as differences between trials with the starting object in the courtyard and trials with the starting object in the building. An additional analysis measured switching costs between the subspaces. For this analysis, we only considered accuracy and RT on non-integration trials that were preceded by other non-integration trials (approximately 25% of all trials). This analysis used a repeated measures 2X2 ANOVA with factors of subspace (building vs. courtyard) and switch (whether the subspace on the preceding trial was the same or different).

#### Free recall task

We tested for subspace effects on object sequential recall probabilities, by using a two-tailed paired-samples t-test to compare the number of consecutively recalled objects that were within the same subspace to the number of consecutively recalled objects that were in different subspaces.

#### Map localization task

Localization errors were computed by assigning each object localization response to the nearest veridical location of an object. If this was the location of the queried object, then the trial was scored as correct. If it was a different object, then the trial was scored as an error. Error trials were then further classified as within-subspace errors (the assigned object was in the same subspace as the queried object) or between-subspace errors (the assigned object and the queried object were in different subspaces). The number of within-subspace errors was compared to the number of between-subspace errors using a two-tailed paired-samples t-test. To test for subspace effects, the number of errors made when the queried object was in the building was compared to the number of errors made when the queried object was in the courtyard using a two-tailed paired-samples t-test.

#### Post-experiment questionnaire

Participants’ rankings of their knowledge of each subspace, use of first-person perspective in each subspace, and use of third-person perspective in each subspace were all compared between the two subspaces (building and courtyard) using two-tailed paired-samples t-tests.

### Functional MRI analysis

#### Estimation of fMRI responses

Voxelwise blood-oxygen level dependent responses during the JRD task were estimated using general linear models (GLMs) implemented in SPM12 (Wellcome Trust Centre for Neuroimaging) and MATLAB (2023a, Mathworks). Trials were assigned to conditions based on the starting object on each trial (which indicated the imagined location), and whether the target object was in the same subspace or different subspace (integration vs. non-integration trials), resulting in 32 total regressors. In addition, a separate GLM was performed in which trials were assigned based on the starting object alone, resulting in 16 regressors. GLMs included regressors for each of the conditions, constructed as impulse functions convolved with a canonical hemodynamic response function. Also included were regressors for the following variables of no interest: head motion parameters (6 regressors), overall framewise displacement of the head (Power et al., 2014), average signals from the CSF and white matter (2 regressors), and differences between scan runs. Temporal autocorrelations were modeled with a first-order autoregressive model. After estimation of the models, the variance-normalized voxelwise responses to each regressor were determined from a t-test contrasting the regressor’s parameter estimate against the resting baseline (Misaki et al., 2010). Functional localizer task runs were analyzed using a similar procedure, with assignment of blocks to four GLM conditions based on the picture category (places, objects, faces, scrambled).

#### Regions of Interest (ROIs)

We defined 5 regions of interest that have been identified in previous work as being involved in spatial memory and/or the perception of environmental scenes. These ROIs were the retrosplenial complex (RSC), parahippocampal place area (PPA), occipital place area (OPA), entorhinal cortex (ERC) and hippocampus (HPC). RSC, PPA, and OPA were defined using group-level masks (parcels) that specified these regions’ locations in 42 participants run in previous lab experiments, following the methodology developed in (Julian et al., 2012). Within these parcels, participant-specific ROIs were defined based on the functional localizer contrast places>objects, by selecting the 200 voxels with the highest contrast value within the corresponding parcel in each hemisphere and then combining across hemispheres to create bilateral ROIs (400 voxels per ROI). The hippocampus and entorhinal cortex ROIs were defined from each participant’s anatomical parcellation as obtained from Freesurfer.

#### Univariate analyses

To test brain activity in the JRD task related to integration (integration vs. non-integration trials) and subspace (building vs. courtyard trials), we first averaged the t-values across all voxels within each ROI for each of the 32 regressors (16 objects x 2 integration conditions), and then averaged these values further across regressors corresponding to the 4 conditions of interest (building+integration, building+non-integration, courtyard+integration, courtyard+non-integration). We then used a 2X2 repeated-measures ANOVA with participant as random effect on these ROI values. To test for relation between these neural effects and corresponding behavioral effects, Pearson correlation was computed between the JRD behavioral integration costs (integration vs. non-integration, for accuracy and RT) and the JRD neural integration costs (integration vs. non-integration in each ROI). Results of all analyses were FDR-corrected across ROIs. Finally, to test for effects across the whole brain, voxelwise SPM contrasts were computed for the main factors (integration vs. non-integration, building vs. courtyard) and corrected for multiple tests using SPM’s multiple comparison correction (random field correction, p=0.05).

#### Multivariate analysis

To explore the representational space within each ROI, we extracted multi-voxel fMRI activation patterns for each participant. These were constructed from the outputs of the 16 or 32 regressor GLMs. Activation patterns for each regressor were created by taking the regressor-specific t-values and then concatenating across all voxels within each ROI. These patterns were then compared to each other using one minus Pearson correlation to create 16x16 or 32x32 representational dissimilarity matrices (RDMs). Coding of spatial location and heading was then tested using these RDMs (see Supplementary Materials for details of heading coding analyses).

We tested for coding of spatial location within each ROI by creating model RDMs where each cell indicated the spatial distance between the corresponding pair of objects (Marchette et al., 2014; Peer & Epstein, 2021). We tested both a global location RDM (which included the distances between all 16 objects) and two subspace-specific location RDMs (which included only the distances between the 8 objects in each subspace). These model RDMs were compared to the neural RDM in each ROI by measuring the Spearman correlation between them. The significance of each location code in each ROI was then estimated using one-tailed one-sample t-tests (with FDR-correction across ROIs).

We then further decomposed the global code model into possible subcomponents. These included the two within-subspace location RDMs, as well as three additional RDMs: an RDM of between-subspace distances (containing only distances between pairs of objects in different subspaces), an RDM of segmentation between the subspaces (1 for between-subspace cells, 0 for within-subspace cells; positive loading corresponds to greater representational dissimilarity between subspaces than within subspaces), and an RDM of scale difference between the subspaces (1 for within-courtyard cells, 0 for within-building cells; positive loading corresponds to greater representational dissimilarity within the courtyard than within the building). We then took the elements from each RDM (within-building, within-courtyard, between-subspaces, segmentation, scale), z-scored each regressor, and performed a linear regression where these regressors are the predictors (plus a constant term) and the neural RDM elements are the result. We then tested each regressor’s beta value for significance across participants in each ROI using one-tailed one-sample t-tests (with FDR-correction across ROIs). Finally, to test if significant regressors could explain the fit to the global location model by themselves, partial correlation was used between each regressor, the neural RDM, and the global location code RDM.

Next, we tested whether integration trials differed from non-integration trials. We constructed a model corresponding to a categorical difference between integration and non-integration trials (1 for different integration condition, 0 for same integration condition integration; positive loading corresponds to greater representational dissimilarity between integration and non-integration trials compared to trials with the same integration status) and compared this to the 32 x 32 neural RDM in each ROI by Spearman correlation. We also separately tested this model for the subset of cells corresponding to representational dissimilarities between building patterns and the subset of cells corresponding to the representational dissimilarities between courtyard patterns (defined, in both cases, by their starting objects). We also performed a decomposition of the neural RDM into subcomponents, using the same five subcomponents described above, defined in this case on the 32 x 32 matrix, plus a subcomponent corresponding to the integration vs. non-integration distinction.

Finally, we performed multidimensional scaling to visualize the representational codes within specific ROIs. Neural similarity matrices were averaged across participants and normalized to the range of zero to one. The cmdscale MATLAB function was used to calculate the multi-dimensional scaling of the data to two-dimensional space.

#### Simulations examining the relationship between univariate and multivariate effects

Finally, we performed simulations to explore the possibility that the observed multivariate effects might arise from differences in univariate activity between conditions. To this end, we simulated fMRI data that mimicked the observed univariate differences but had no underlying multivariate code, then tested whether effects emerged in the multivariate analyses.

The simulations had several assumptions. First, we assumed that all conditions shared the same underlying multivariate pattern, i.e. some voxels are consistently more active than others, across all conditions. Second, we assumed that univariate activity differences between trials manifested as a multiplication of activity by a factor that was the same in all voxels: for example, if trial 1 activates an ROI twice as much as trial 2, then every voxel in the ROI will be twice as active in trial 1 than in trial 2. This assumption was based on the idea that the underlying multivariate pattern is caused by voxels having different numbers of neurons responding to the task, and univariate differences are caused by a proportionate increase in firing that applies equally to all neurons. Thus, for example, a univariate difference between conditions might be caused by every neuron increasing its firing rate by 50%, which would lead to voxelwise response increases that would be proportionately the same across voxels (assuming a linear transform between aggregate neural activity with a voxel and its BOLD response). Finally, we assumed that there was an equivalent level of noise (either measurement or trial-related noise) for all conditions.

Our hypothesis was that—if the assumptions above were true—then conditions with higher univariate response would have a higher signal-to-noise ratio than conditions with lower univariate response. Multivariate activation patterns for the higher-response conditions would then be more similar to each other than multivariate activation patterns for the lower-response conditions. To test this idea, we created “simulated ROIs” for each participant corresponding to each of our real ROIs. In each simulated ROI, we selected a random baseline underlying multivariate pattern. We then simulated response patterns in each condition, by taking each participant’s actual ratios of activity between conditions in the ROI (building+non-integration, building+integration, courtyard+non-integration, courtyard+integration) and multiplying our simulated patterns by these differences. We then generated 8 different patterns for each condition (corresponding to 8 “objects”) by taking the corresponding pattern (after multiplication according to the condition) and adding random noise. Finally, we created simulated neural RDMs by correlating the simulated patterns to each other, and we performed the RDM regression described in the previous section between the model RDMs (within-subspace distances, between-subspace distances, etc.) and the simulated neural RDM. We then tested the fit to the different regressors by one-tailed one-sample t-tests.

## RESULTS

### Environmental learning

To explore the neurocognitive mechanisms underlying the coding of hierarchical spaces, we familiarized participants with a virtual environment consisting of a courtyard containing a single building that could be entered through a doorway on its “West” side (Fig. 1). Thus, the environment was divided into two physically connected but visually separated subspaces (courtyard, building interior) that were organized in a hierarchical fashion (building contained within courtyard). Within the environment were 16 objects whose locations were learned by the participants—eight along the boundaries of the courtyard and eight along the interior walls of the building.

We implemented a multi-stage Environmental Learning task to ensure that participants were familiarized with the layout of the environment and the locations of the objects within it. Initially, all objects were visible, but as training progressed, increasing numbers of the objects were covered by wooden boxes to induce reliance on spatial memory (see Materials and Methods). By the end of the learning procedure, all participants could successfully navigate to all object locations without making more than one error, even when all the objects were covered (Fig. S1). The Environmental Learning task took 27 min on average (range: 14–49 min). On day 2, participants repeated the learning task from the beginning and exhibited near-perfect accuracy (average of 99.4% correct object localizations).

### Behavioral evidence for segmentation into subspaces

To understand the mental representation that participants formed of the virtual environment, we examined behavioral performance during the Environmental Learning task and in two post-learning memory tasks (map localization and free recall). The first question we asked was whether participants formed a unitary, fully integrateds cognitive map of the whole environment, or if instead they mentally segmented it into two separate spatial parts for the two subspaces (i.e., building and courtyard). By design, our paradigm was intended to induce segmentation into subspaces: the courtyard and the building interior were perceptually distinct and visually separated from each other, with orthogonal principal axes; moreover, object learning was organized by subspaces in the first six stages of the learning task. Results from three analyses suggested that participants did indeed form representations that reflected this subspace division.

First, we analyzed participants’ errors in the last two stages of the learning task (Fig. 2A). In these stages, participants were required to navigate by memory to object locations that could either be in the building or the courtyard. Almost all errors (88.5%) were mis-localization of objects within the correct subspace— that is, going to an incorrect location within the building when the target location was in the building, or going to an incorrect location in the courtyard when the target location was in the courtyard. Almost no errors involved going to locations in the wrong subspace. The difference between within-subspace and between-subspace errors was significant (p=0.009, two-tailed paired-samples t-test).

**Figure 2:**
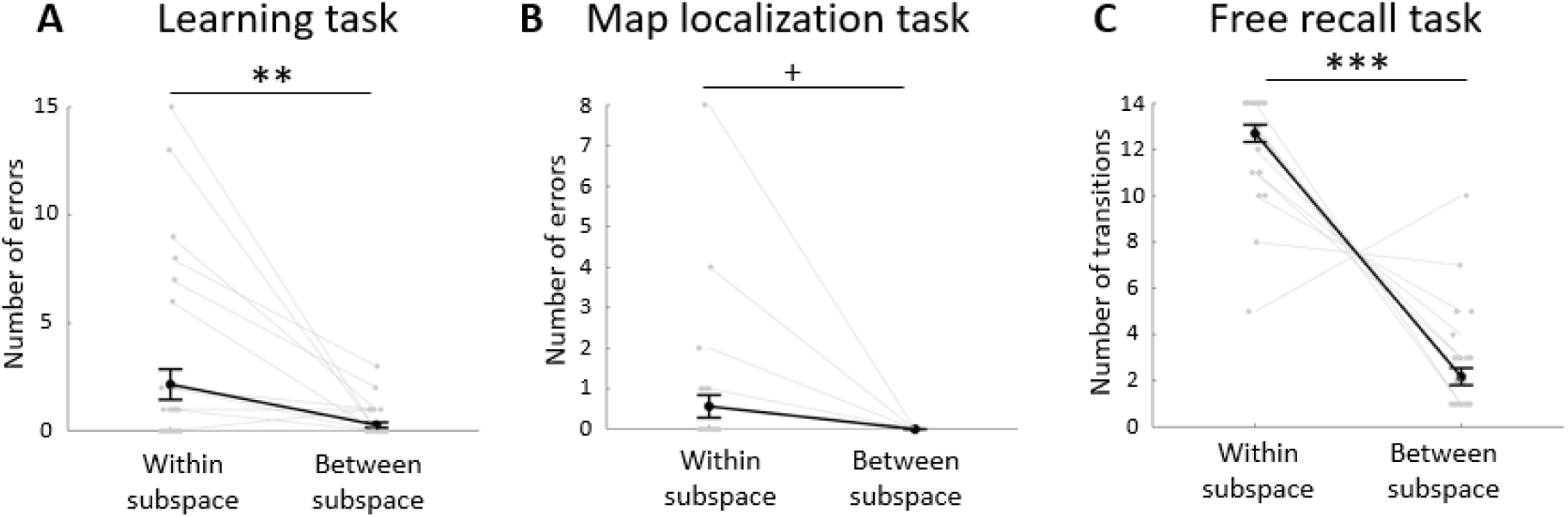
Behavioral evidence for segmentation into subspaces. A) In the environmental learning task, participants made more within-subspace errors than between-subspace errors. B) A similar trend was observed in the map localization task. C) During free recall, within-subspace transitions between consecutively named objects were more frequent that between-subspace transitions. Grey points and lines indicate individual subjects, black points and lines indicate the group average. Error bars indicate standard error of the mean. * - p<0.05, ** - p<0.01, *** - p<0.001, + - marginal effect (0.05<p<0.1).

Second, we examined errors in the map localization task (Fig. 2B). Similar to the pattern observed during learning, objects were sometimes mis-localized within the correct subspace, but they were never mis-localized into the wrong subspace (0.6 and 0 errors on average respectively, marginal effect at p=0.053).

Finally, we looked at transitions between objects in the free recall task (Fig. 2C). Consecutively recalled objects were more often in the same subspace than in different subspaces (12.7 vs. 2.2 consecutive recalls, respectively, p<0.0001), indicating that participants did not recall objects randomly, but instead organized their free recall based on subspaces. Indeed, the majority of participants (20 out of 32) recalled all the objects in one subspace and then all the objects in the other subspace, and therefore had only one between-subspace transition. Of these, 17 started their recall in the courtyard and completed in the building whereas 3 started their recall in the building and completed it in the courtyard. The remaining 12 participants had more than one recall transition between the subspaces; in these participants, there was no bias toward transitions in either direction (p=0.72, two-tailed paired-samples t-test).

Together, these behavioral results suggest that participants did not form a unitary, fully-integrated representation extending across the building and the courtyard in an equipotential manner. Rather, they mentally segmented the environment into two subspaces. This division into subspaces was evident in spatial learning, spatial recall, and free recall.

### Behavioral evidence for integration across subspaces

We next explored how participants integrated the building and the courtyard representations into a larger whole. In other words, how did they succeed in understanding the spatial relationships between locations in different subspaces, given that the two subspaces were represented—to some extent—separately? To address this question, we analyzed behavioral responses in the JRD task. On each trial, participants imagined themselves standing in front of one object (the starting object) and reported the relative location (left vs. right) of a second object (the target object). For half of the trials, the starting and target objects were in different subspaces (i.e., one object in the building and the other in the courtyard), while for the other half, both objects were in the same subspace (Fig. 3A). Thus, the first kind of trial required integration across the two subspaces, while the second kind of trial did not require integration. We reasoned that comparison between these two trial types might reveal behavioral signatures of a spatial integration mechanism that operated on integration (between-subspace) trials but not on non-integration (within-subspace) trials.

**Figure 3:**
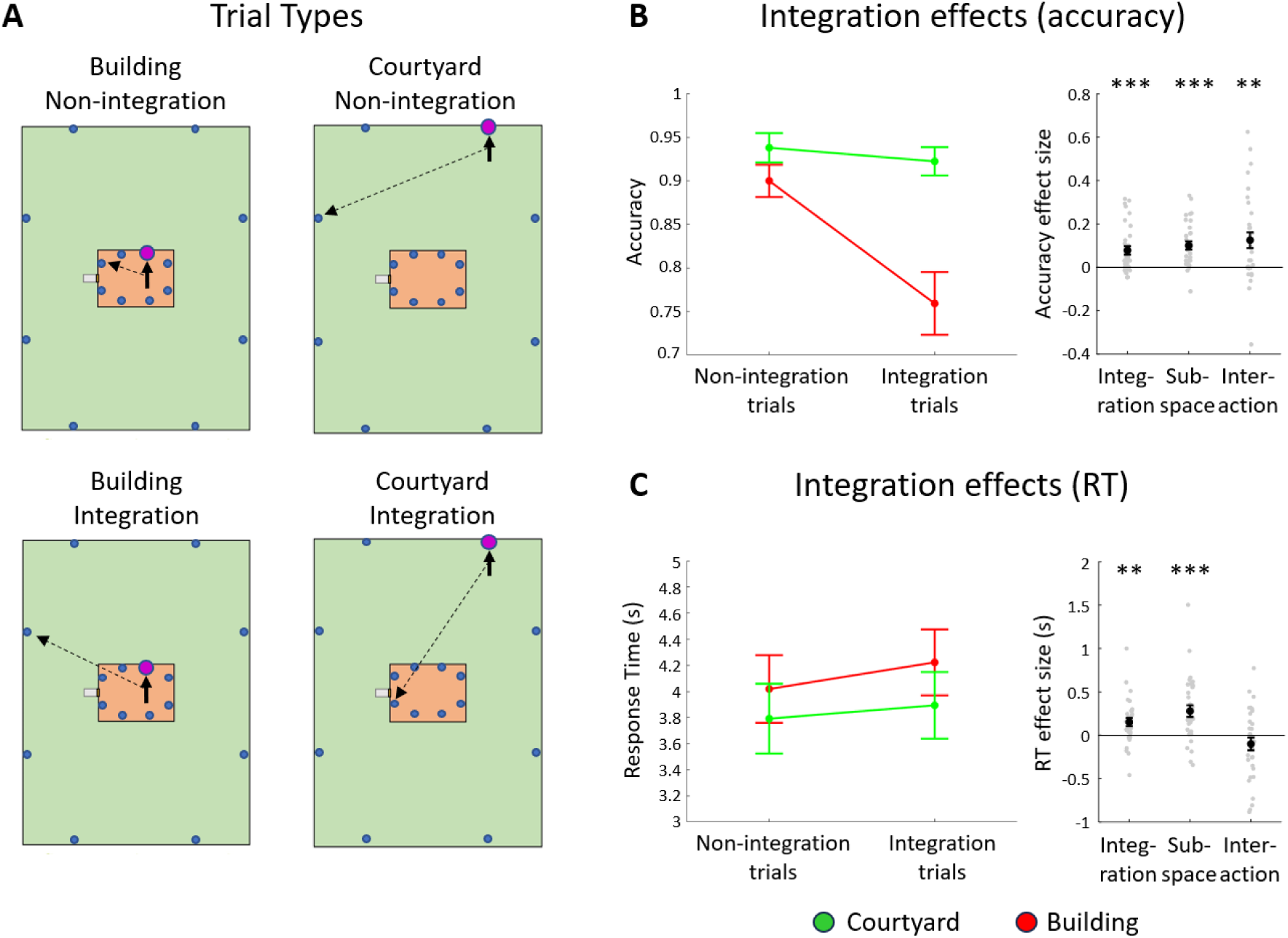
Behavioral evidence for integration between the two subspaces. A) Example schematics of non-integration (within-subspace) and integration (between-subspace) JRD trials; the red dot indicates the starting object, the full arrow indicates the facing direction, and the dashed arrow indicates the target object’s direction. B) Integration costs: integration trials show lower accuracy, with an interaction with starting subspace demonstrating asymmetric responses. This is consistent with a model in which people use the hierarchical relations to infer the object direction when they are in the higher hierarchy layer (courtyard) by pointing to the location of the building instead of the location of the object in it. Left – mean+-SE for each trial type. Right – effect sizes: integration – integration cost, non-integration minus integration trial accuracy. Subspace – subspace effect – courtyard minus building accuracy. Interaction – interaction between effects – (courtyard-non-integration minus courtyard-integration) minus (building-non-integration minus building-integration). C) Integration costs in response times (RTs). Integration trials show higher RT than non-integration trials, and building trials show higher RT than courtyard trials, but there is no interaction. Elements similar to panel B, but effect size calculations reversed (integration minus non-integration and building minus courtyard). Other plot elements similar to Figure 2.

To test this idea, we performed 2x2 ANOVAs on response times (RT) and accuracies, with the main factors being integration (integration vs. non-integration) and the location of the starting object (building vs. courtyard; Fig. 3A). The results showed a clear integration effect: integration trials had lower accuracy than non-integration trials (accuracy=84%,92%, respectively, F=15.6, p=0.0004; Fig. 3B) and longer response times (RT=4.1,3.9, respectively, F=10.8, p=0.003; Fig. 3C). Thus, there was an integration cost that was evident in both accuracy and response time. This suggests that integration trials engaged an extra cognitive process that was not engaged on non-integration trials.

Notably, the integration cost on accuracy (but not RT) was asymmetric across the subspaces. The ANOVA on accuracy revealed a significant interaction between integration and starting subspace (F=11.8, p=0.002), reflecting the fact that the integration penalty was significant for starting objects in the building (paired t-test, p=0.0005) but not for starting objects in the courtyard (paired t-test, p=0.22). This asymmetry suggests that participants may have used their knowledge of the nested structure of the environment when making their responses. Specifically, because they knew that all targets in the building were contained inside the building, they could respond accurately on courtyard-integration trials by thinking about the position of the building within the courtyard without taking the further step of integrating the building interior space into the courtyard space. In contrast, they could not make the inference in the opposite direction, so building-integration trials could only be solved by integrating the two subspaces, and an integration penalty was observed. The ANOVA on RT did not reveal a similar interaction between integration and starting subspace (ANOVA F=1.96, p=0.17); for this measure, the integration cost was found for both building trials (paired t-test p=0.02) and courtyard trials (paired t-test marginal effect of p=0.08).

The main factor of location (building vs. courtyard) was also significant in the ANOVA: trials with the starting object in the building had lower accuracy (accuracy=83%,93%, respectively, F=29.0, p=0.000007) and longer response times (RT=4.1,3.8, respectively, F=17.0, p=0.0003) than trials with the starting object in the courtyard. This echoes a similar effect that was observed in the Environmental Learning task: participants made more localization errors on day 1 for building objects than for courtyard objects, (4.8 vs. 2.6 on average, p=0.006, two-tailed paired-samples t-test; Fig. S2A), and they also reported that their knowledge of locations was worse for building objects that for courtyard objects in post-learning questionnaires (average ranking=9.2, 8.5, respectively, p=0.003; Fig. S2B). These effects may relate to the fact that the courtyard was larger than the building: the average separation between the objects was 282 virtual meters in the courtyard vs. 67 virtual meters in the building. Consequently, participants spent more time in the courtyard than in the building during learning (63% vs. 37%), giving them more time to encode the object locations, with greater temporal separation between the objects which may have made them more distinct. Alternatively, these effects may relate to differences in imagery between the subspaces, as participants reported they imagined the courtyard more from a third-person perspective and the building marginally more from a first-person perspective (p=0.004,0.06, respectively; Fig. S2B).

Finally, we also analyzed the JRD task in terms of switching costs between the subspaces. Previous work used switching cost as evidence for representational distinction between hierarchical levels (Brockmole & Wang, 2002), and we wished to test for a similar effect. We restricted our analyses to responses on non-integration trials, because these trials can be completed by thinking about one subspace alone without consideration of the other subspace. We tested whether accuracy and RT on these trials were affected by whether the locations accessed in the immediately preceding trial (in cases where this trial was also a non-integration trial) were in the opposite subspace (switch) or same subspace (no switch) (Fig. 4A). A 2x2 ANOVA with factors of subspace (courtyard vs. building) and switch vs. no-switch found no significant effects on accuracy (ps>0.28; Fig. 4B). However, there was a significant switching cost in RT (F=18.6, p=0.0002), which interacted with subspace (F=5.75, p=0.02; Fig. 4C). Specifically, the switching cost was observed for building-to-courtyard transitions (relative to courtyard-to-courtyard transitions; paired t-test p=0.00006), but it was not observed for courtyard-to-building transitions (relative to building-to-building transitions; paired t-test p=0.53). Thus, we observed an asymmetry between the subspaces – switching from the building to the courtyard incurs a penalty, but switching from the courtyard to the building does not. The main effect of subspace was not significant (F=0.59, p=0.45). These results support the idea that the courtyard and the building are representationally distinct, insofar as a switching cost is observed. Moreover, the asymmetric nature of the switching cost is generally consonant with the fact that subspaces are hierarchically related to each other, making transitions upwards (building to courtyard) different from transitions downwards (courtyard to building).

**Figure 4:**
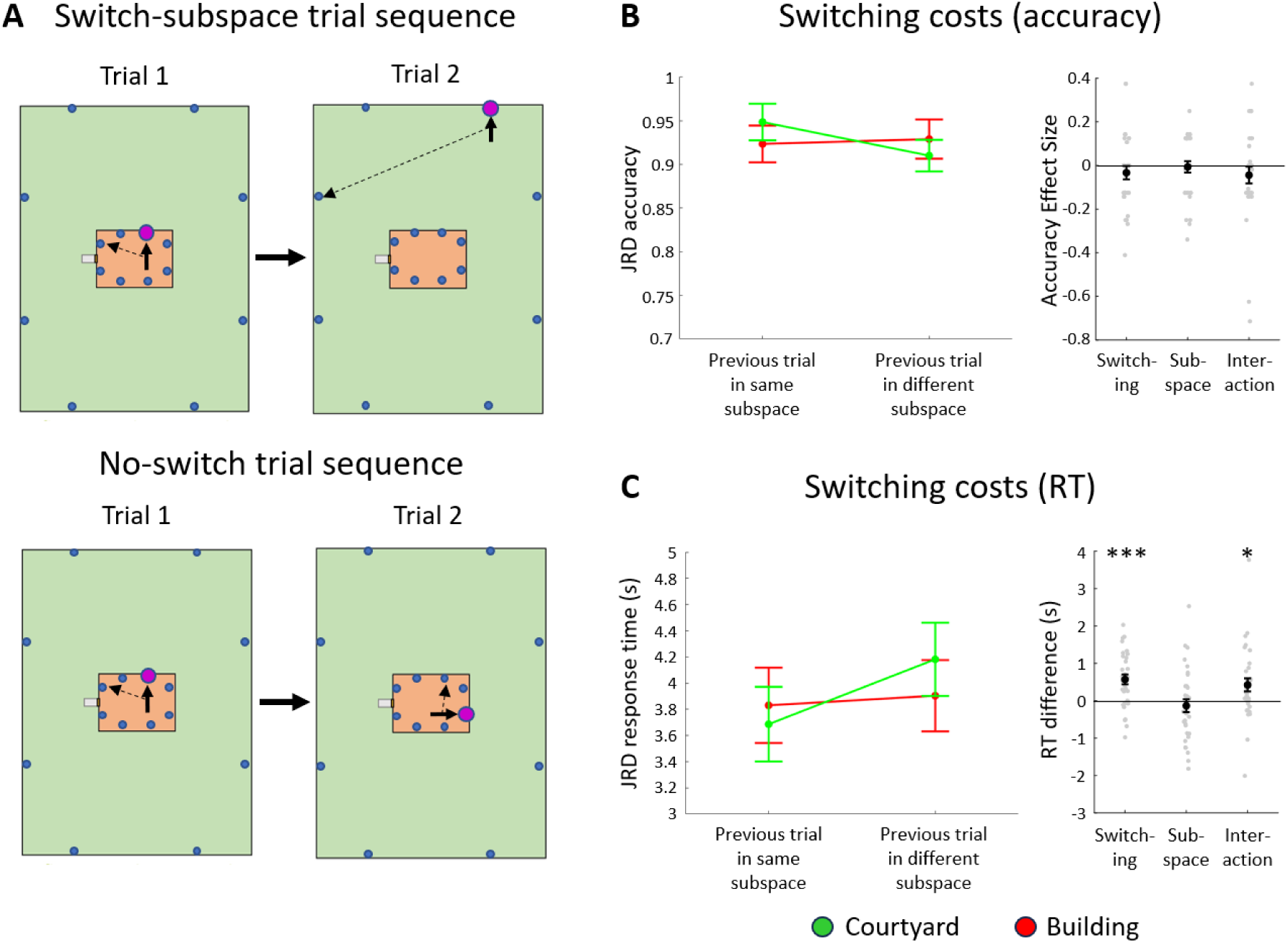
Behavioral switching cost effects between the two subspaces. A) Example schematics of switch and no-switch JRD trial pairs; the red dot indicates the starting object, the full arrow indicates the facing direction, and the dashed arrow indicates the target object’s direction. B) Switching costs in accuracy – no significant effects. Switching – switching costs – same-subspace minus different-subspace; Subspace – subspace effect – courtyard minus building accuracy. Interaction – interaction between effects – (courtyard-no-switch minus courtyard-switch) minus (building-no-switch minus building-switch). C) Switching costs in response time – data shows a switching cost suggestive of asymmetrical representation, such that consecutive trials in a different starting subspaces take more time than in the same subspace, and an interaction shows that this effect is more pronounced in building-to-courtyard transitions than in courtyard-to-building transitions. Elements similar to panel B, but effect size calculations reversed (different minus same subspace, and building minus courtyard). Other plot elements similar to Figure 3.

### Neural evidence for an integration mechanism

The behavioral data described in the preceding sections suggests that participants represented the environment as two separate subspaces that they integrated when required using an active process. To identify the neural locus of this process, we examined fMRI activity during the JRD task. As in our behavioral analyses, we divided trials into 4 conditions, based on the 2x2 crossed factors of integration (integration vs. non-integration) and subspace of the starting object (building vs. courtyard). We then examined univariate responses across these 4 trial types. We focused our analyses on five brain regions that have been identified in previous work as being involved in spatial memory and/or the perception of environmental scenes. These ROIs were the retrosplenial complex (RSC), parahippocampal place area (PPA), occipital place area (OPA), entorhinal cortex (ERC) and hippocampus (HPC).

Our primary hypothesis was that brain areas involved in integration would exhibit different levels of response for integration vs. non-integration trials. Analysis of variance found such differences (i.e., main effect of integration) in RSC, PPA, OPA, and ERC (Fs=12.5, 8.4, 17.7, 6.9; ps=0.001, 0.007, 0.0002, 0.01) with a marginal effect in HPC (F=4.1, p=0.052). Notably, the integration effect did not manifest in the same way in all these regions (Figs. 5A, S3). In RSC and PPA, activity was higher for integration trials than for non-integration trials. In OPA, the opposite pattern was observed: activity was *lower* for integration trials than for non-integration trials. Finally, the two medial temporal lobe regions – ERC and HPC – displayed a third pattern: like OPA, activity was lower for integration trials than for non-integration trials; however, responses to both conditions were below the resting baseline, so this pattern might be interpreted as greater *deactivation* for integration trials than for non-integration trials.

**Figure 5:**
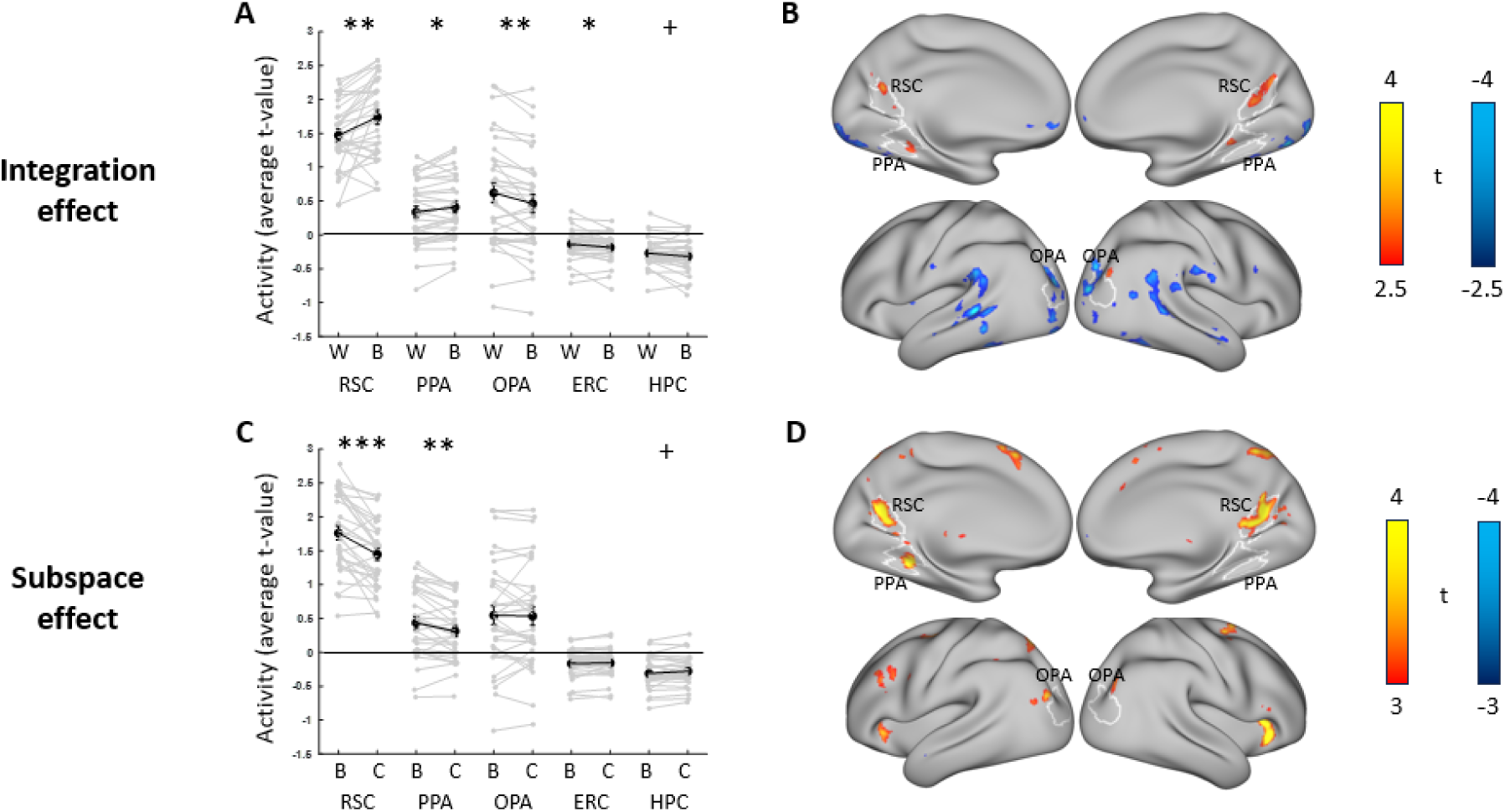
Integration and subspace effects in univariate fMRI response. A) Neural activity related to integration (within-subspace trials (W) that do not require across-subspace integration, vs. between-subspace trials (B) that require integration). The RSC and PPA show increased activity during trials that require between-subspace integration; OPA shows reduced activity during between-subspace integration trials; and ERC and HPC show deactivation during both trial types, and this deactivation is stronger in between-subspace integration trials. B) Whole-brain analysis (uncorrected) shows that the increased activity during between-subspace integration trials is localized to RSC, PPA and a region anterior to OPA, while integration-induced deactivations occur across multiple parts of the lateral occipito-temporal cortex. C) Activity in ROIs during trials in which the imagined location is in the building (B) and trials when it is in the courtyard (C). The RSC and PPA show increased activity during building trials compared to courtyard trials. D) Whole-brain analysis (uncorrected) shows that the increased activation during building trials is localized to RSC, PPA, a region anterior to OPA, and additional brain regions mainly in the insula and prefrontal cortex. Plot elements similar to Figure 2.

A whole-brain analysis of the integration vs. non-integration main effect found no significant voxels after correction for multiple comparisons across the entire brain. However, when the threshold was lowered (to uncorrected p=0.01), we observed patterns that were broadly consistent with the ROI findings (Fig. 5B)—stronger activity in integration trials in RSC, PPA and an additional region immediately anterior to right OPA, and stronger activity in non-integration trials in several lateral occipito-temporal cortex regions, including OPA. Overall, these results show that all spatial system ROIs exhibit integration effects and that these effects are mostly specific to these spatial system regions.

The 2x2 ANOVAs also revealed main effects of subspace (courtyard vs. building) in RSC and PPA (Figs. 5C, S3). Both regions were more active during building trials than during courtyard trials (p=0.00007,0.002, respectively). No other ROI exhibited a significant subspace effect (ps>0.05). In a whole-brain analysis, no voxels exhibited significant subspace effects after correction for multiple comparison, but uncorrected results showed higher activity for building trials than for courtyard trials in RSC and PPA, consistent with the ROI results, and also in a region anterior to OPA, with additional clusters in the medial superior parietal lobe, medial and lateral prefrontal cortex, and anterior insula (Fig. 5D). There was no interaction between the integration and subspace effect in any ROI (all ps>0.35) and no significant voxels for the interaction in a whole-brain FDR-corrected analysis.

Thus, we find that when a trial requires integration of the two subspaces, activity increases in RSC and PPA and decreases in the other ROIs. Can any of these neural effects be linked to the behavioral effects of integration? To examine this, we looked at whether individual differences in the neural integration effects correlate with individual differences in the behavioral integration effects observed in accuracy and RT. We found no correlations with the accuracy integration effect, although there were marginal negative trends in OPA (r=-0.41, p=0.07) and ERC (r=-0.39, p=0.07). For the RT integration effect, on the other hand (Fig. 6), we observed a significant positive relationship in RSC (r=0.5, p=0.008) and significant negative relationships in OPA (r=-0.47, p=0.01) and HPC (r=-0.64, p=0.003), with marginal trends in PPA (r=0.31, p=0.08) and ERC (r=-0.34, p=0.07). Thus, participants with larger behavioral integration effects in RT exhibited greater positive neural integration effects (integration>non-integration) in RSC and greater negative integration effects (non-integration>integration) in OPA and HPC.

**Figure 6:**
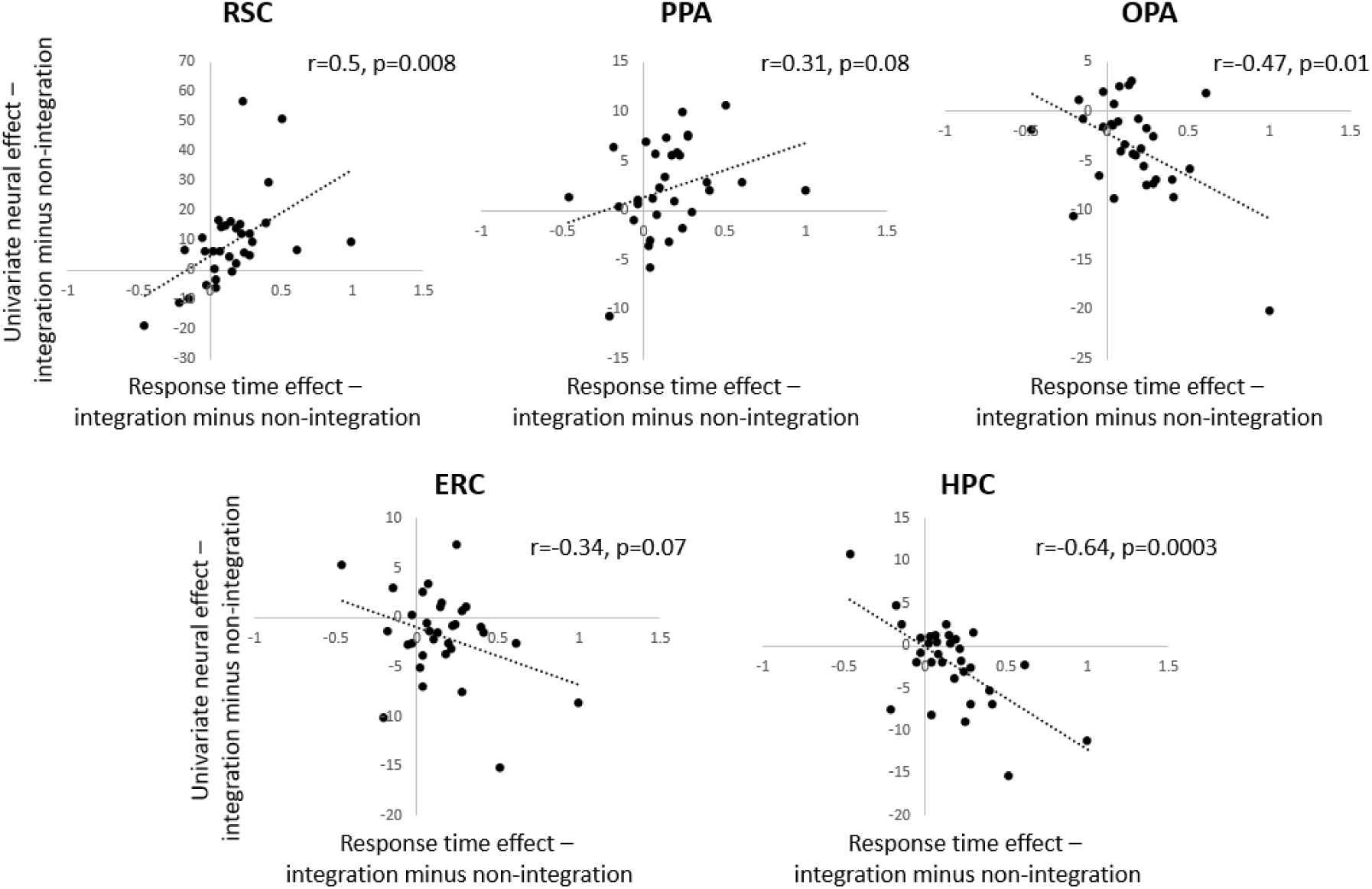
Correlation between neural and behavioral effects. Dots represent individual subject values, dashed lines indicate regression line fit.

In sum, our results show neural integration effects in three different groups of brain regions. The RSC, PPA (and potentially a region anterior to OPA) are more active for trials requiring integration across the two subspaces. These regions are also more active during building trials than during courtyard trials. A second group of regions, hippocampus and entorhinal cortex, are deactivated during the task, and this deactivation is stronger when integration across subspaces is required. Finally, OPA is more active during non-integration trials than during integration trials. The neural integration effects in RSC, OPA, and HPC were most closely related to the behavioral integration effects, which suggests that these regions are the most likely candidates for the neural locus of the spatial integration mechanism. We consider possible explanations for these effects in the Discussion.

### Multivoxel pattern analyses reveal representations of subspace segmentation and subspace scale

We next turned to the question of how the spatial structure of the virtual environment was represented in the brains of our participants. To this end, we performed representational similarity analyses on the multivoxel activation patterns evoked during the JRD task. In previous studies, we found evidence that similarities and differences between these activation pattern could be explained by spatial quantities that vary across the trials, such as the imagined facing direction (heading) and imagined location (Marchette et al., 2014; Peer & Epstein, 2021; Vass & Epstein, 2013). We applied the same methodology here. First, we constructed neural similarity matrices for each ROI by calculating the pairwise similarities between the multivoxel patterns evoked by the 16 possible starting objects, which correspond to 16 different imagined locations (with corresponding headings) within the virtual environment. Then we compared these to representational similarity matrices corresponding to similarities in imagined headings and imagined location.

Surprisingly, given our findings in previous studies (Marchette et al. 2014), we found no evidence for coding of imagined heading (See Supp. Materials). In contrast, we did find strong evidence for coding of imagined location. First, we tested a global location model that included Euclidean (straight-line) distances between all 16 objects (Fig. 7A-B). There was significant correlation to this model in RSC (p=0.0002), PPA (p=0.002) and HPC (p=0.006) and marginal correlation in OPA and ERC (both ps=0.053). Next, we tested subspace-specific location models, which only included distances between the 8 objects in each subspace. Here we observed only marginal coding in OPA of the distances between the building objects (p=0.06)— all other within-building distance codes were not significant (all ps>0.28) and no area encoded within-courtyard distances (all ps>0.71).

**Figure 7:**
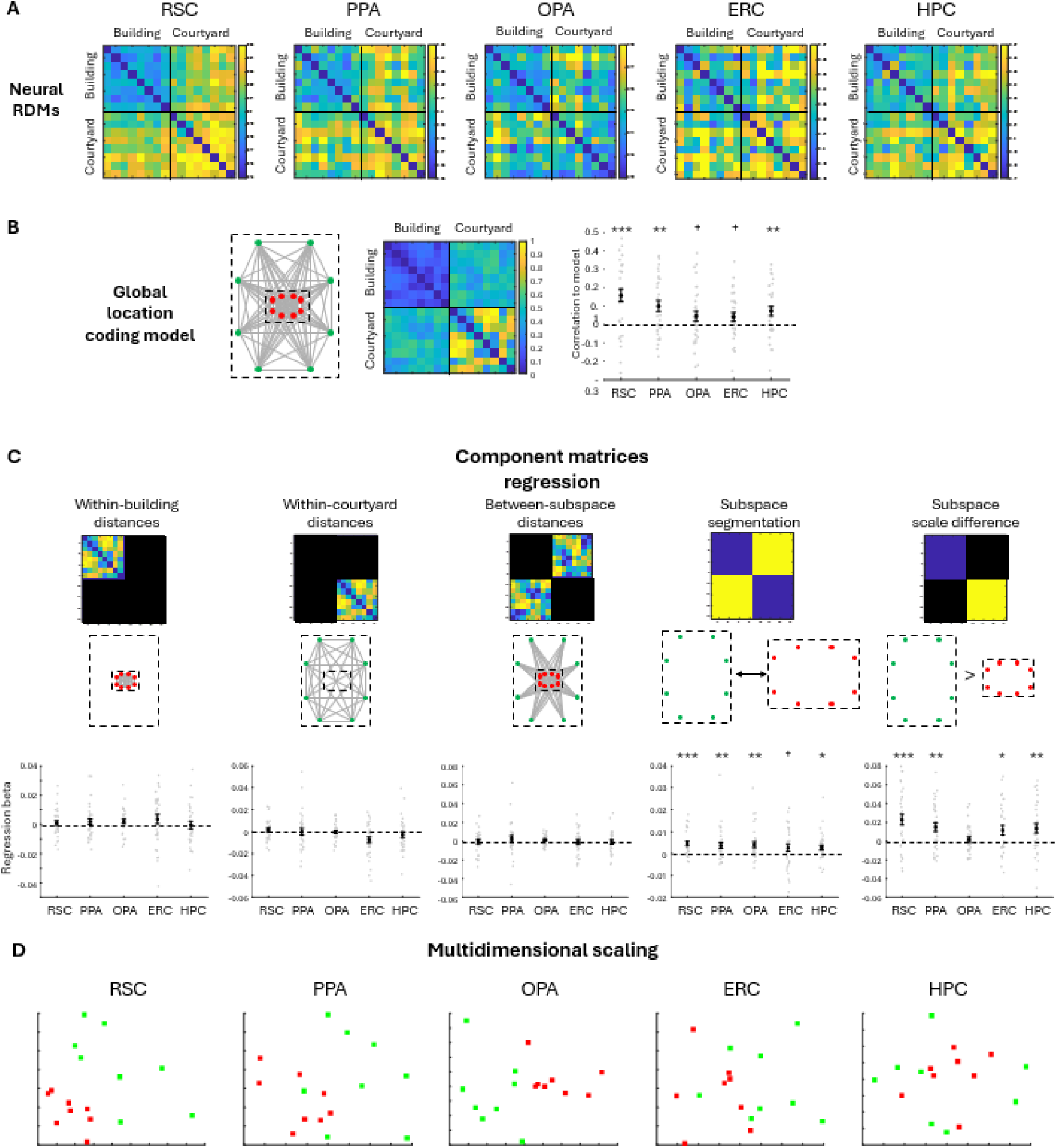
Multivariate patterns show segmentation and scale differences between the subspaces. A) Multivariate pattern dissimilarity between neural patterns corresponding to the 16 objects in each ROI. B) Left – a scheme of the environment with all objects’ distances; middle – the expected pattern dissimilarity if neural pattern distance corresponds to inter-object pattern distance; Right – correlation of the veridical distances model to the neural dissimilarity matrices across ROIs. All ROIs showed marginal or significant fit to the neural model. C) A regression of different factors that could together create the fit to the veridical distance model: distance variability between building objects, distance variability between courtyard objects, distance variability between building and courtyard objects, segmentation between the building and courtyard, and scale difference between the building and courtyard. The segmentation and scale effects are significant in most ROIs, but the inter-object distances within and between subspaces are not. Plot elements same as Figure 2. D) Multidimensional scaling of the pattern correlations in each ROIs (red – patterns corresponding to building object trials, green – patterns corresponding to courtyard object trials). In most ROIs, patterns for building objects are visibly separate than those of courtyard objects, and there is a difference in scale (building patterns are more clustered / similar to each other).

What causes this fit to the global location code model, in the absence of within-subspace distance coding? To explore this issue further, we regressed the neural similarity matrix for each brain region against a model containing five factors (Fig 7C): 1) distances between objects in the building, 2) distances between objects in the courtyard, 3) distances between objects in different subspaces, 4) categorical segmentation into two subspaces, and 5) difference in scale between the two subspaces. We found no significant loading on the first three regressors (all ps>0.3, across ROIs). However, the segmentation effect was significant in RSC, PPA, OPA and HPC and marginal in ERC (p=0.0004, 0.008, 0.006, 0.01, 0.07, respectively), indicating that there was a categorical difference between the building and courtyard patterns. In addition, the scale effect was significant in RSC, PPA, ERC and HPC (p=0.0009,0.003,0.02,0.008, respectively; Fig. 7C), demonstrating that building patterns were more closely clustered (had less neural distance between them) than courtyard patterns.

We then tested whether the segmentation and scale effects could explain the fit of the global location model. Using partial correlation, we found that the global location model remained significant when accounting for the segmentation effect in RSC, PPA and HPC (all ps<0.01) and remained marginal in ERC (p=0.07), but was no longer significant in these ROIs when accounting for the scale effect (all ps>0.16). In the OPA, the global location model was significant when accounting for the scale effect (p=0.048), but was no longer significant when accounting for the segmentation effect (p=0.15). Thus, the neural similarity patterns that we observe can be explained by segmentation between the subspaces in OPA and scale differences between the subspaces in RSC, PPA, ERC and HPC.

Finally, we performed multidimensional scaling on the pattern similarity matrices to visually examine the segmentation and scale effects (Fig. 7D). Across the five ROIs, a separation between the building and courtyard objects could be observed, as well as a more condensed clustering of the building objects compared to the courtyard objects. These results visually demonstrate the existence of the segmentation and scale effects. Overall, our results demonstrate a categorical effect of representational segmentation between the courtyard and the building and a representational scale difference between them.

### Multivoxel pattern analyses reveal that spatial representations are affected by spatial integration

The previous analyses examined spatial coding of the imagined location (defined by the starting object) without consideration of how these codes might be affected by the integration condition (defined by the relationship between the starting object and the target object). To address this issue, we reanalyzed the data, this time separately defining integration and non-integration activation patterns for each starting object (32 patterns total; Fig 8A). We then repeated the regression analysis of the previous section, but with a sixth regressor added to account for the categorical separation between integration and non-integration trials (Fig. 8B).

**Figure 8:**
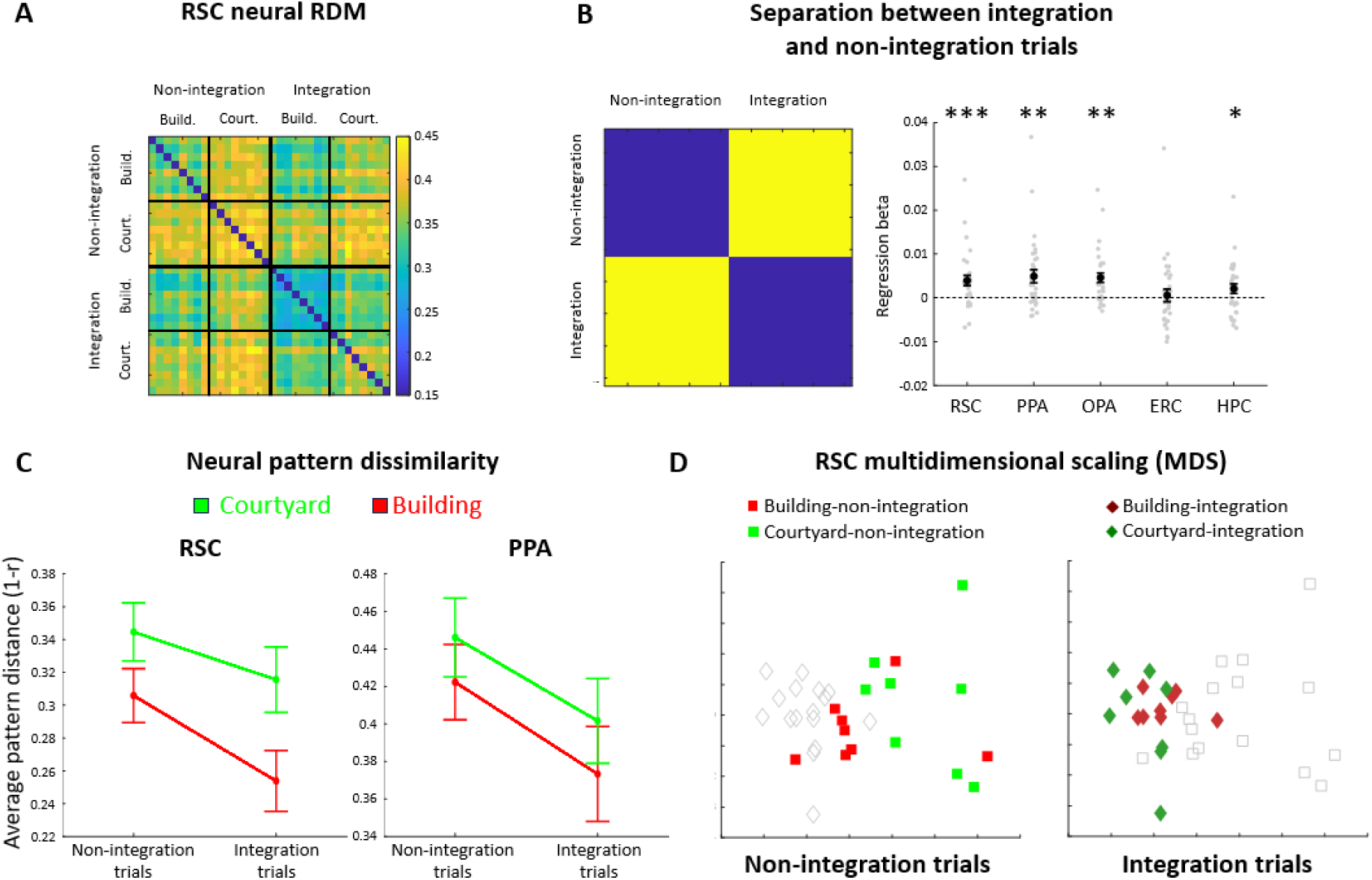
Multivariate patterns show separation and scale differences between integration and non-integration trials. A) Multivariate pattern dissimilarity between neural patterns corresponding to the 32 regressors (16 objects, separated to between-subspace – integration – and within-subspace – non-integration – trials) in RSC. B) Regression results for the separation between integration and non-integration trials in each ROI. Left – model matrix, right – fit to the model matrix in each ROI. Plot elements same as Figure 2. C) Neural pattern dissimilarities across conditions in RSC and PPA. Multivoxel activity is dissimilar between subspaces and between integration and non-integration trials. D) Multidimensional scaling of the pattern correlations in RSC. Patterns for between-subspace (integration) trials are separate and more clustered than within-subspace (non-integration) trials. In addition, within each condition (integration and non-integration), patterns for building objects are separate and more clustered than those of courtyard objects.

This analysis revealed a significant fit to the integration regressor in RSC, PPA, OPA and HPC (ps=0.001,0.001,0.001,0.02; Fig. 8B), indicating that activation patterns on integration trials differed from activation patterns on non-integration trials. Further analyses revealed that this difference between integration and non-integration patterns was observed when the starting object was in the building (ps=0.002,0.001,0.005,0.03) and also when the starting object was the courtyard (ps=0.0005,0.006,0.006,0.04). The integration factor was not significant in ERC (overall, p=0.25; building trials p=0.11; courtyard trials p=0.73; Fig. 8B).

These results show that the spatial codes in RSC, PPA, OPA, and HPC are modified when participants are required to integrate. But how are they changed? To investigate this, we first examined whether the scale of the representation (the average representational dissimilarity between patterns) differed between integration and non-integration trials. Fig. 8C shows the average pattern dissimilarity between the building objects and the courtyard objects under both integration conditions in RSC and PPA. In both regions, integration trials were less dissimilar to each other (i.e., more clustered) than non-integration trials (RSC p=0.002, PPA p=0.0006). In OPA, ERC and HPC this effect was not observed (ps=0.37,0.34,0.18). Greater clustering of integration trials compared to non-integration trials could also be observed by visual inspection after multidimensional scaling (Fig. 8D).

### Relation between the multivariate and univariate effects

The results above indicate an interesting parallel between the univariate and multivariate effects. In several of our ROIs, building trials evoke greater activation than courtyard trials, and they also have greater pattern similarity. Similarly, integration trials evoke greater activation and have greater pattern similarity than non-integration trials. We therefore explored whether the higher multivariate pattern similarity within these trial types could be explained by the higher univariate activation.

We specifically considered a scenario in which there was a single underlying response pattern in each ROI across all trial types. If this were the case, then we hypothesized this pattern would be less affected by noise on trial types with higher activation, and hence the multivariate activity patterns for these trial types would be more similar. We performed a simulation to test this idea. For each ROI, we constructed 32 simulated activity patterns corresponding to the 32 conditions in our data (8 objects in the building and courtyard trials, with separate patterns for integration vs. non-integration trials), by starting with the same fixed baseline pattern (chosen at random, with positive values) for all 32 conditions. We then scaled activity across conditions in the same ratio as the univariate differences observed in this ROI in our experimental data, and then added random “measurement noise” to each pattern. Thus, the average “voxelwise” values of the patterns differed across conditions, but the scale of the measurement noise was the same. We then explored the multivariate pattern correlations within and across trials.

We found that the simulation resulted in multivariate effects that mirrored the patterns observed in the real data (Fig. 9). First, all simulated ROIs demonstrated a separation between the subspaces, with the ERC demonstrating the weakest effect (p=0.005, 0.002, 0.002, 0.01, 0.002, for RSC, PPA, OPA, ERC and HPC respectively). Second, the simulated RSC and PPA demonstrated a scale difference between the building and the courtyard, and the HPC demonstrated a marginal effect, but no such effect was observed in OPA or ERC (p=0.001, 0.004, 0.06, 0.80, 0.64). Third, all simulated ROIS showed a separation between non-integration and integration trials (p=0.02,0.006,0.007,0.007,0.002; although in the actual fMRI data this effect was not significant in ERC).

**Figure 9.**
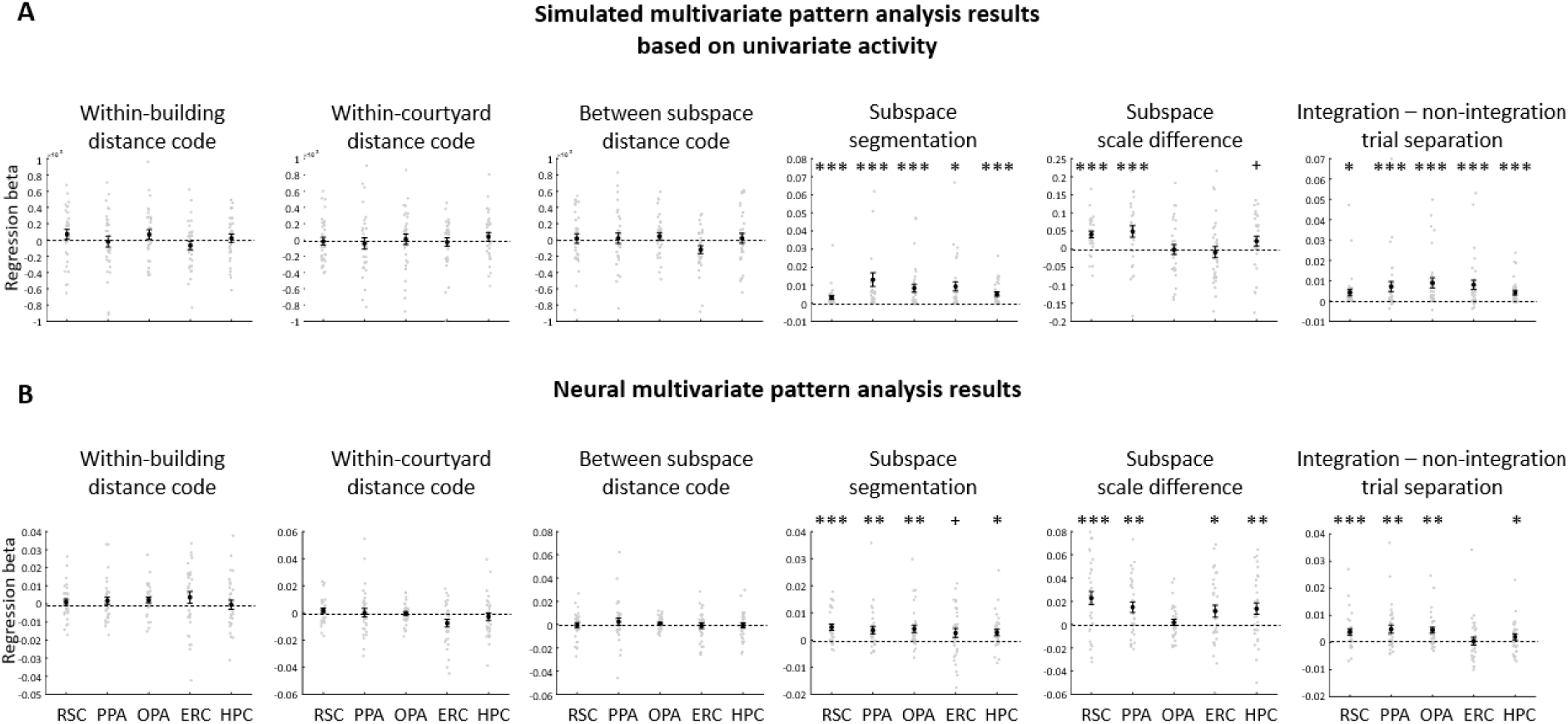
Simulated multivariate pattern analysis results. A) Simulated results, based on univariate activity differences between conditions (see main text). B) The real experimental results. Both the simulated and real results show similar effects (subspace separation, subspace scale difference, integration vs. non-integration trial separation), suggesting that the observed multivariate results could potentially be explained by the univariate differences between experimental conditions. Plot elements similar to Fig. 2.

These simulation results suggest that the observed multivariate effects could, in theory, arise from univariate differences between the conditions, even in the absence of true neural differences between the multivariate voxelwise activation patterns. We consider the implications of this finding below in the Discussion.

## DISCUSSION

Many of the environments we encounter in our everyday lives are hierarchically organized – they are composed of smaller spaces nested within larger spaces. The goal of this study was to understand how our mind/brains form cognitive maps of such nested spaces. To this end, we trained participants on a virtual environment consisting of a building inside a courtyard, and we scanned them with fMRI while they performed a JRD task that required them to think about spatial relationships within and across the subspaces. We found that trials requiring integration across the subspaces had decreased accuracy and longer reaction times compared to trials that did not require integration. These behavioral differences were accompanied by univariate fMRI response differences in scene-responsive and medial temporal lobe brain regions, which were correlated with the behavioral effects in RSC, OPA, and HPC. Multivoxel pattern analyses revealed spatial representations that reflected the hierarchical organization of the environment, with evidence for segmentation into subspaces (RSC, PPA, OPA, and HPC), preservation of the relative spatial scales of the subspaces (RSC, PPA, ERC, and HPC), and modulation of the spatial representations by spatial integration (RSC and PPA). However, our simulations indicated that these multivariate effects could potentially be attributed to the univariate activity differences between task conditions. Overall, these results indicate that people form cognitive maps of nested environments that reflect their hierarchical structure and then use an active cognitive process when required to integrate across the levels of the hierarchy.

Our findings build on previous work suggesting that segmentation into subspaces is a common feature of cognitive map organization. Adjacent spaces with boundaries between them tend to be represented as separate, as indicated by both behavioral results (Han & Becker, 2014; Marchette et al., 2017; McNamara, 1986; Newcombe & Liben, 1982; Wiener & Mallot, 2003) and neural findings (Jeffery, 2024; Kim & Maguire, 2018; Marchette et al., 2014; Peer & Epstein, 2021). We observed evidence for segmentation in several aspects of our data. In the environmental learning and map localization tasks, participants made within-subspace errors than across-subspace errors. During free recall of the objects in the environment, the order of recall reflected the subspace organization—indeed, the majority of our participants recalled all the objects in one subspace before transitioning to the other. In the JRD task, integration and switching costs were observed between the subspaces, replicating previous findings (Bilge & Taylor, 2010; Brockmole & Wang, 2002; Wang & Brockmole, 2003b, 2003a). Together these results indicate that participants represented the environment by mentally segmenting it into two separate subspaces.

Why might hierarchical environments be represented as segmented, instead of as a global map that extends across subspaces and thus is intrinsically integrated? One advantage of segmentation is that it reduces the computational cost of navigational planning. There are fewer objects to maintain in working memory when navigating within each subspace, making it beneficial to chunk the items by subspace in memory (Brockmole & Wang, 2005). This can be especially true when people spend prolonged periods of time within each subspace and only occasionally move between them, and when there are few connection points between the subspaces (in our case, a single door). Therefore, segmentation of space into separate hierarchical levels / spatial scales may be part of a general mechanism of organizing space to reduce the cognitive burden of navigation.

The disadvantage of segmented representations for hierarchical environments is that it can be challenging to make between-level spatial judgments and navigational decisions (e.g. planning how to get from a location in a building to another location in the city). Our behavioral results suggest that people meet this challenge by applying an integration mechanism that has an additional computational cost, as evidenced by a response time increases and accuracy decreases. One interesting aspect of this integration mechanism is that this process appears to be asymmetrical – participants were more accurate making top-down inferences (courtyard to building) than bottom-up inferences (building to courtyard). According to participants’ reports, this was due to their use of knowledge of the hierarchical structure to make top-down inferences by judging directions to the building containing the object instead of to the object itself. This illuminates one of the main advantages of having an hierarchical representation – it can simplify between-subspace judgments and top-down hierarchical planning (Brockmole & Wang, 2005; Wiener & Mallot, 2003).

Our fMRI results provide insight as to how this integration mechanism is implemented in the brain. We found that several brain regions exhibited univariate response differences on the JRD task between trials that required participants to think about the spatial relationship between the hierarchical levels and trials that did not require integration across subspaces. Specifically, RSC and PPA exhibited increased activity on integration trials, whereas OPA, ERC, and HPC exhibited decreased activity. The RSC finding is consistent with previous work that showing that this region is involved in understanding the spatial relationships between locations that are separated from each other (Epstein et al., 2007; Peer et al., 2019). The PPA finding is more surprising, insofar this region is often assumed to be primarily involved in the representation of the immediately surrounding space (Epstein et al., 2007). However, we note the anterior parts of the PPA have been shown to be involved in analyzing environments that are larger than a single scene (Peer et al., 2019), and anterior PPA, RSC, and a region anterior to OPA have been shown to be functionally connected (Baldassano et al., 2016; Silson et al., 2016) into an anterior scene network that has been implicated in scene memory (Steel et al., 2021). Notably, our whole-brain analyses revealed a region immediately anterior to the OPA that was more activated in integration trials than in non-integration trials, suggesting that this entire anterior network may be involved in spatial integration. The reduced response in OPA during integration trials is consistent with its putative role in scene perception (Bonner & Epstein, 2017; Kamps et al., 2016) and processing of local (within-subspace) spatial relationships (Peer & Epstein, 2021). The reduced response in ERC and HPC took the form of deactivation during all trials, but greater deactivation during integration. This might be either interpreted as relating to their involvement in the default-mode system which deactivates during tasks that require concentration (Raichle, 2015), or alternatively this might be an actual activation increase at the neuronal lever that presents as BOLD signal deactivation (Ekstrom, 2021; Hill et al., 2021). Crucially, these fMRI integration effects were correlated with behavioral integration effects, with the strongest correlations in RSC, OPA, and HPC. Together, these results indicate that these brain regions support an active mechanism that allows people to integrate between levels of a hierarchical space when such integration is required.

One unexpected finding in our study was that the two subspaces differed in terms of their behavioral and neural responses: participants had slower and less accurate responses for spatial judgments when their imagined location (as indicated by the starting object) was in the building compared to the courtyard, and fMRI responses in RSC and PPA were greater for building trials than for courtyard trials. The RT and fMRI effects did not show an interaction with the integration effects (i.e., they were not affected by the location of the target object), indicating that they had an independent genesis. These subspace effects may relate to differences during learning. In the courtyard, the objects were more spatially separated than in the building. Consequently, participants spent more time during learning in the courtyard and the courtyard objects were more temporally separated. Participants also started each learning stage in the courtyard, even stages where the objects to be found were in the building. All these factors may have led them to form a better representation of the courtyard than the building. Indeed, participants made fewer errors in the courtyard during learning, and they self-reported that their knowledge of locations in the courtyard was better than their knowledge of locations in the building. Thus, the longer RT and greater activation for building trial in the JRD task may reflect a greater memory retrieval load on these trials. Alternatively, these effects may be more directly related to the size of the spaces. Independent of how well the spaces were learned, participants may have found it more effortful to form spatial images of smaller spaces during the JRD task, and RSC and PPA responses might be greater when imagining spaces that are more closely bounded (mirroring a similar effect in scene perception; (Henderson et al., 2011)). Overall, it is unclear what caused the differences between the building and the courtyard, and further studies are needed to understand what is driving these effects.

Our multivoxel pattern analyses revealed several effects that were related to the hierarchical organization of the environment. First, we observed a segmentation effect: locations in the two subspaces were representationally distinct from each other. Second, we observed a scale effect: locations in the courtyard were more representationally distinct than locations in the building. Third, we observed effects related to integration: integration trials were representationally distinct from non-integration trials, and integration trials were more representationally clustered than non-integration trials. These effects were localized to the same regions that showed univariate activity differences. The consilience of these findings with the behavioral results is notable. However, we are reluctant to draw strong conclusions from them because our computational simulation suggested that—if certain assumptions hold, the multivariate effects could be a byproduct of univariate differences. Skepticism about the multivariate effects may be further warranted by the fact we were not able to identify any within-subspace distance or heading codes, in contrast to our previous studies of similar environments (Marchette et al., 2014; Peer & Epstein, 2021).

Finally, our findings may have implications for understanding the role of hierarchy in memory organization beyond space. Hierarchical organization exists in many domains; for example, language (words, sentences, paragraphs), semantic memory (nested categories), and working memory (chunking). Two potential benefits exist to hierarchical organization. First, by organizing items hierarchically, the number of items in each level is reduced, enabling their maintenance and manipulation in working memory. This may enable visualizing all items together and understanding the relations between each pair of items, a process which becomes exponentially harder as the number of items increases (Brockmole & Wang, 2005; Dirlam, 1972; Johnson, 1970; Mandler, 1967, 2013). In our study, hierarchical organization may have served to facilitate memory of locations and spatial relations within each subspace, thus simplifying within-subspace spatial judgments. Second, organizing memory hierarchically enables the use of the hierarchical structure to make inferences about item relations, as evidenced in hierarchical planning where planning is performed separately for different scales/hierarchy levels (Tomov et al., 2020; Wiener & Mallot, 2003). In our study, participants appear to have used hierarchical inference to make correct spatial judgments by pointing toward the building instead of toward the specific items within the buildings. We speculate that our findings regarding the brain systems involved in the coding of hierarchical spatial environments may be relevant to memory organization in other domains, especially given the evidence that these systems play a role in representing both spatial and non-spatial cognitive maps (Behrens et al., 2018; Bellmund et al., 2018; Epstein et al., 2017; Peer et al., 2015). Although our results highlight the role of medial temporal lobe and scene regions in making simple hierarchical judgments in the spatial domain, other brain regions, such as frontal lobe regions, may become involved when hierarchical planning demands become more abstract or more complex (Balaguer et al., 2016).

In conclusion, our findings show that people represent each level of hierarchically organized environments separately, and use an active mental process mediated by scene-responsive and medial temporal lobe brain regions to integrate across the levels. This study adds to the increasing literature demonstrating that cognitive maps—at least, in some circumstances—are not unitary spatial fields, but instead are structured entities that involve the representation of distinct spatial subspaces and the spatial relationships among those subspaces. These findings can be relevant to understanding how memory in general is organized by segmentation and integration processes.

## Author contributions

Conceptualization, investigation, analysis, visualization, writing – MP+RAE. Software development – MP.

