## Supplementary Materials for "Cognitive maps for hierarchical spaces in the human brain"

### Investigation of heading codes

To test whether neural patterns in each ROI represent the imagined heading when facing an object, we tested heading models of local and global heading coding using representational similarity analysis (RSA; (Kriegeskorte, 2008; Marchette et al., 2014)). To test for global heading coding, we created a model RDM where each cell indicated if the imagined heading between the corresponding pair of objects is the same or different. We also created another model RDM where each cell indicated the absolute angular difference between the two imagined headings. To test for local heading coding, we created matrices (for each subspace) by taking the corresponding subset of the global heading RDM that reflects only the heading differences between objects belonging to that subspace. Each of these model RDMs was compared to the neural RDM in each ROI, by measuring the Spearman correlation between them. The significance of each heading code in each ROI was then estimated using one-tailed one-sample t-tests (with FDR-correction across ROIs).

We tested for heading codes in two ways. First, we tested for a global representation of heading across the entire environment (i.e. “facing North” evokes the same pattern in both the building and the courtyard). No ROI demonstrated significant coding of global heading; i.e. objects with the same global heading do not evoke patterns that are more similar than objects with different global headings (marginal heading code in OPA, $p=0.057$; all other ROIS $ps>0.13$). We next looked at coding of local heading within each subspace separately, since our previous work suggests that heading codes in different subspaces can use reference frames that are globally inconsistent with each other—particularly in environments like the current one where the primary axes of the subspaces are misaligned. No local heading codes were observed in either subspace in any ROI (all $ps>0.25$). Results were similar when neural patterns were analyzed in terms of angular similarity between headings (all $ps>0.05$; marginal heading code in the building in OPA and ERC, and in the courtyard in RSC, $p=0.09, 0.07, 0.07$, respectively).

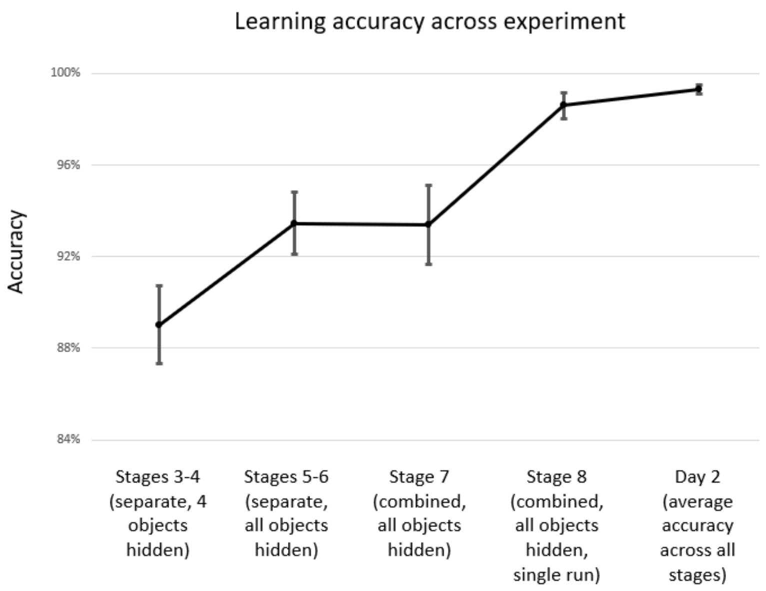


**Figure S1: Learning performance.** Performance improved throughout learning, reaching near-perfect performance on day 2 (0.3 errors on average across all day 2 learning stages). Data from stages 1-2 are not shown because all objects were visible and therefore errors are not informative. Error bars indicate standard error of the mean.

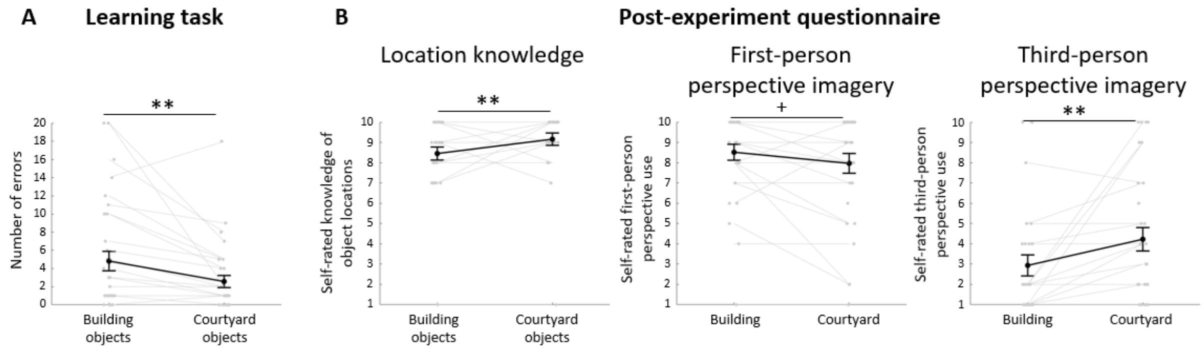


**Figure S2: Differences between the building and courtyard.** A) Participants made more learning errors in the building than the courtyard. B) Participants rated their knowledge of the building as worse than of the courtyard, and reported more use of first-person perspective imagery and less use of third-person perspective imagery in the building compared to the courtyard. Plot elements similar to Figure 2.

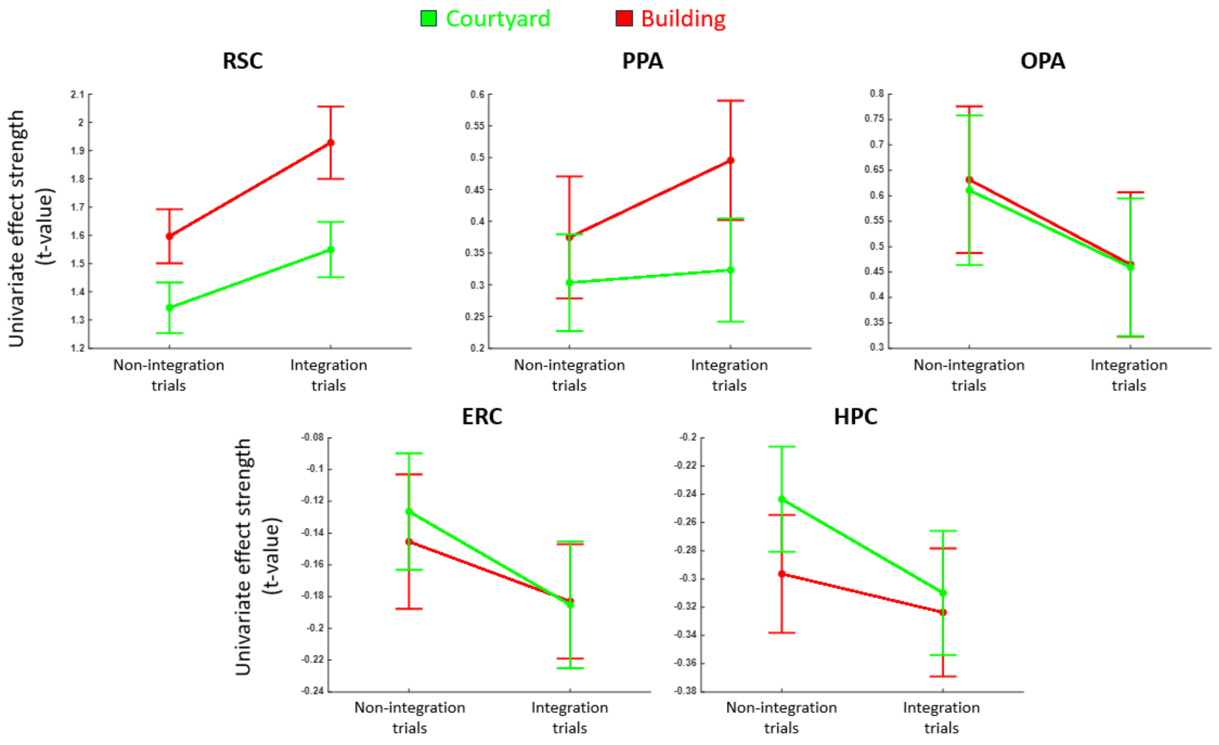


**Figure S3: Activity profiles for the different trial types in each ROI.** Figure elements similar to Fig. 2.
